# Reinvigoration of translational activity in dysfunctional T cells initiates the early intratumoral response to PD-1 blockade

**DOI:** 10.1101/2025.09.24.676875

**Authors:** Paulien Kaptein, Nadine Slingerland, Anne M. van der Leun, Eline Runderkamp, Roos A. Wagensveld, S. Michael Chin, Janniek R. Mors, Mercedes Machuca-Ostos, Timm M. Reissig, Kelly D. Moynihan, Ivana M. Djuretic, Yik A. Yeung, Ton N. M. Schumacher, Daniela S. Thommen

**Affiliations:** Division of Molecular Oncology and Immunology, Oncode Institute, The Netherlands Cancer Institute, Amsterdam, The Netherlands; Asher Biotherapeutics, Inc., South San Francisco, California; Division of Tumor Biology and Immunology, The Netherlands Cancer Institute, Amsterdam, The Netherlands; Department of Hematology, Leiden University Medical Center, Leiden, The Netherlands

**Author notes:** Department of Cancer Biology, Dana-Farber Cancer Institute, Boston, MA, USA. These authors contributed equally to this work. These authors share senior authorship (Ton N. M. Schumacher, Daniela S. Thommen).

## Abstract

T cells are key effectors of antitumor responses elicited by PD-1 blockade. However, it remains elusive by which mechanism(s) PD-1 blockade initiates T cell-driven antitumor immunity in cancer tissues. Here, we dissect early T cell reactivation upon anti-PD-1 in patient-derived tumor fragments. Using bispecific antibodies to target anti-PD-1 to individual T cell subsets, we demonstrate that intratumoral CD8^+^ and CD4^+^ T cells can independently drive immune remodeling of the tumor microenvironment. The CD8^+^ and CD4^+^ T cells that respond to anti-PD-1 exhibit a shared dysfunctional gene program, characterized by tumor-reactivity, terminal exhaustion, effector capacity, and reduced translational activity. Notably, rather than acting through transcriptional rewiring, anti-PD-1 reinvigorates dysfunctional T cells by overcoming this translational barrier, resulting in restored effector function. Altogether, these results reveal dysfunctional T cells as initiators of early tissue responses to PD-1 blockade and identify a novel mode of their therapeutic reinvigoration through restoration of translational control.

**One Sentence Summary:** Translational reactivation of tumor-residing dysfunctional T cells drives early intratumoral immune activity upon PD-1 blockade.

## INTRODUCTION

Tumor-specific T cells play a pivotal role in antitumor immunity, but intratumoral T cell function is profoundly inhibited by negative signals within the tumor microenvironment (TME) (*1, 2*). In particular, chronic exposure to tumor antigens in the TME drives T cells to differentiate into a state of dysfunction, also called exhaustion (*3–5*). Dysfunctional T cells are characterized by a gradual reduction of proliferative capacity, impairment of effector function, and elevation of inhibitory receptor expression such as PD-1, CTLA-4, and LAG-3 (*4, 6–8*). Therapeutic blockade of these receptors, referred to as immune checkpoint blockade (ICB), can boost antitumor T cell function. In particular, blockade of the PD-1/PD-L1 axis has demonstrated remarkable clinical success (*9–12*), and anti-PD-1/anti-PD-L1 treatment has become a cornerstone in the treatment of multiple types of cancer (*13*). Nevertheless, despite its profound clinical impact, the cellular and molecular mechanisms by which PD-1/PD-L1 blockade mediates antitumor immunity remain incompletely understood.

Recent studies have identified a critical role of T cells primed or activated in the lymph node for durable responses to PD-1 blockade, either via replenishment or replacement of the tumor-specific T cell pool in tumor tissues (*14–17*). In addition, emerging evidence suggests that anti-PD-1 also reinvigorates T cells that already reside at the tumor site, which may act synergistically with the effects of PD-1 blockade in lymph nodes (*18, 19*). However, a number of aspects of PD-1 blockade-induced T cell activation in the TME remains unresolved. First, the relative role of the intratumoral CD8^+^ and CD4^+^ T cell compartments for the initiation of the local immunological response following PD-1 blockade has not been established. CD8^+^ T cells are considered main effector cells of anti-PD-1 therapy due to their direct cytotoxic capacity (*20–22*). Moreover, CD8^+^ T cell counts serve as a strong prognostic marker across many cancers (*23*). In contrast, the specific role of CD4^+^ T cells in the response to PD-1 blockade remains less well understood. Notably, early genetic ablation studies revealed that PD-1 restrains both CD8⁺ and CD4⁺ T cells (*24, 25*), and accumulating evidence supports a significant role of CD4^+^ T cells in cancer immunity (*26–28*). CD4^+^ and CD8^+^ T cell subsets have both been shown to harbor tumor-reactive T cell receptors (TCRs) (*29, 30*), express high levels of exhaustion markers within the TME (*31–33*), be capable of tumor cell killing (*34*) and respond to checkpoint blockade (*18, 35, 36*). Furthermore, these subsets also share the ability to produce key cytokines (*37*), such as interferon-gamma (IFN-γ), which enhances antigen presentation, inhibits tumor cell growth, and induces chemoattractants, thereby facilitating the recruitment of additional immune cells to the TME (*38*).

Next to the lack of clarity on the relative role of CD4^+^ and CD8^+^ T cells in intratumoral immune reactivation upon PD-1 blockade, uncertainty exist around the differentiation state of the responding T cells. In mouse models, the T cell pool required for durable antitumor immunity has been shown to exhibit a "precursor-like" phenotype with stem-like potential and high proliferative capacity, characterized by intermediate PD-1 levels and expression of the transcription factor *Tcf7* (*14, 15, 39, 40*). These cells, however, predominantly reside in lymphoid tissues. Hence, the specific state of T cell subsets responding at the tumor site remains undefined. Upon entering the human TME and (re-)exposure to tumor antigen, tumor-specific T cells rapidly undergo terminal differentiation towards TCF7^−^ TOX^+^ dysfunctional cells expressing exhaustion markers such as CD39 and CXCL13 (*4, 8, 41–43*). Given their high levels of PD-1 (*41*), intratumoral dysfunctional T cells seem potential targets for PD-1 blockade therapy. In further support of this concept, tumor reactivity is enriched within the dysfunctional pool (*29, 44*), and its presence in pre-treatment biopsies was found predictive for therapeutic outcome following PD-1 blockade in cancer patients (*41, 45–47*). At the same time, it has been difficult to reconcile these observations with the low reinvigoration potential of this subset as observed in preclinical models, likely because of irreversible epigenetic changes acquired by cells prior to establishment of the dysfunction-associated gene expression program (*48–51*).

To investigate T cell reinvigoration following PD-1 blockade in human cancer tissues, we employed a previously established patient-derived tumor fragment (PDTF) model that (1) preserves the spectrum of dysfunctional T cell states ex vivo, (2) captures the earliest immunological changes induced by anti-PD-1, and (3) makes it possible to analyze the functional consequences of defined perturbations on T cells within their native tissue context (*18, 52*). Importantly, prior work has established a tight correlation between immune reactivation induced by PD-1 blockade in this ex vivo model and treatment response of the same patient (*18*), underscoring the physiological relevance of this model. By combining single-cell protein and transcriptomic analyses with mechanistic experiments in PDTFs, we here characterize the phenotypic, transcriptomic, and functional changes in T cells and the human TME within hours to days after ex vivo PD-1 blockade. Furthermore, to disentangle the contribution of CD8^+^ and CD4^+^ T cells to PD-1 blockade response, we employ novel bispecific antibodies to selectively direct PD-1 blockade to either the CD8 or CD4 lineage. Our results demonstrate that both CD8^+^ and CD4^+^ T cells can independently drive downstream cytokine and chemokine responses upon PD-1 blockade, but with varying contributions across tumors. The reinvigorated T cell subsets exhibit a highly dysfunctional (or exhausted) state at baseline, with a shared gene program between CD8^+^ and CD4^+^ subsets. Mechanistically, PD-1 blockade reactivated dysfunctional T cells at the tumor site by restoring effector function predominantly at the posttranscriptional/translational level, and with minimal transcriptomic changes. Notably, the reactivated T cell pool displays enhanced protein translation capacity, pointing towards a novel molecular mechanism of reinvigoration of this subset. Collectively, these results demonstrate that tumor-residing dysfunctional CD8^+^ and CD4^+^ T cells are independent key initiators of the intratumoral immune response to PD-1 blockade, challenging previous notions regarding the inert nature of these cell pools and emphasizing their therapeutic significance.

## RESULTS

### T cell-targeted anti-PD-1 bispecific antibodies selectively induce PD-1 signaling in CD4^+^ or CD8^+^ T cells

Prior work has established that ex vivo PD-1 blockade enhances the production of proinflammatory soluble mediators in PDTFs for a subset of human tumor samples, and that such increased production strongly correlates with clinical outcome to PD-1 blockade (*18*). Furthermore, this TME remodeling was shown to be dependent on intratumoral T cells (*18*). In line with prior observations (*18*), we found that in tumors in which PD-1 blockade resulted in enhanced secretion of soluble mediators not only intratumoral CD8^+^ T cells, but also CD4^+^ T cells acquire increased expression of the CD137 and OX40 activation markers (**Fig. 1, A and B)**. With the aim to understand the individual contributions of CD4^+^ and CD8^+^ T cells to TME remodeling, we developed bispecific antibodies that simultaneously target PD-1 and CD4 (αCD4xαPD1) or PD-1 and CD8 (αCD8xαPD1), respectively. The specificity of these bispecific antibodies was confirmed by co-staining with a competing αPD-1 antibody (**Fig. 1C**). Moreover, titration of αCD4xαPD1, αCD8xαPD1, and a clinically used αPD-1 antibody (nivolumab) confirmed that the bispecific antibodies specifically blocked PD-1 staining on their designated T cell subsets, with an IC50 that is comparable to that of nivolumab (**Fig. 1D**).

**Figure 1.**
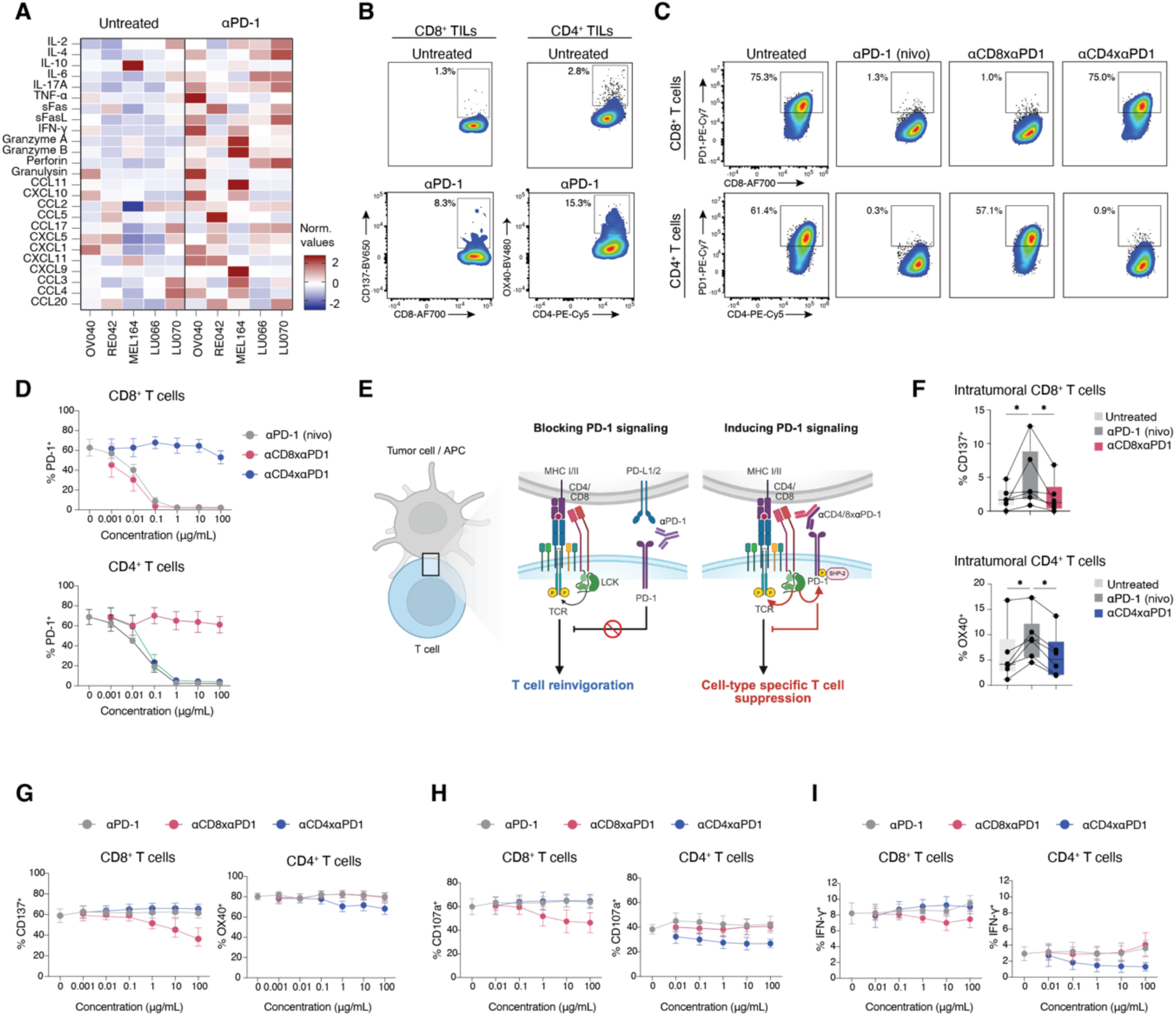
Bispecific αPD1xαCD4 and αPD1xαCD8 molecules allow cell type-specific PD- 1 inhibition. (**A**) Heatmap displaying z-scores of normalized values of 25 soluble mediators secreted by untreated and anti-PD-1-treated PDTFs (n = 5). (**B**) Representative flow cytometry plots of CD8^+^ (CD137) and CD4^+^ (OX40) T cell activation upon ex vivo anti-PD-1 treatment of PDTFs. (**C**) Representative flow cytometry plots of PD-1 staining of CD4^+^ and CD8^+^ T cells from PBMCs that were stimulated for 48 hrs with αCD3 + αCD28 in the presence of either αPD-1, αCD8xαPD1 or αCD4xαPD1. (**D**) PD-1 staining of CD4^+^ and CD8^+^ T cells from PBMCs that were cultured in the presence of αCD3 + αCD28 combined with the indicated concentrations of either αPD-1, αCD8xαPD1, or αCD4xαPD1 (n = 5) for 48 hrs measured by flow cytometry. (**E**) Schematic overview of mechanism of action of αPD-1, leading to T cell reinvigoration, and proposed mechanism of action of αCD8xαPD1 and αCD4xαPD1, leading to specific T cell inhibition. (**F**) Quantification of CD137 on total intratumoral CD8^+^ T cells and OX40 on total intratumoral CD4^+^ T cells from untreated, αPD-1, αCD8xαPD1, and αCD4xαPD1-treated PDTFs, as measured by flow cytometry (n = 6). \**P* < 0.05 by Friedman test corrected for multiple comparisons. (**G-I**) Flow cytometry analysis of the activation status of CD4^+^ and CD8^+^ T cells from PBMCs that were stimulated with αCD3 + αCD28 in the presence of the indicated concentrations of either αPD-1, αCD8xαPD1, or αCD4xαPD1 for 48 hrs. Activation status was measured by analysis of CD137 expression for CD8^+^ T cells and OX40 expression for CD4^+^ T cells (**G**), the degranulation marker CD107a (**H**), and the effector cytokine IFN-γ (**I**) (n = 3).

Next, we aimed to assess the functional activity of the bispecifics. A priori, the mode of action of the bispecific antibodies is difficult to predict. On the one hand, these bispecific molecules could serve as regular antagonists of PD-1/PD-L1 signaling, with the added specificity of targeting either CD4^+^ or CD8^+^ T cells. Alternatively, the bispecifics might function as cell-specific PD-1 agonists. Specifically, both the CD4 and CD8 co-receptors are intimately associated with the tyrosine kinase Lck, which is essential for initiation of the TCR signaling cascade upon antigen engagement (*53*). The bispecifics may therefore bring Lck into close proximity to PD-1, subsequently promoting downstream PD-1 signaling in the respective subset (**Fig. 1E)**, a mechanism reminiscent of the previously described PD-1 signaling inhibition via enforced recruitment of the CD45 phosphatase (*54*). In support of the latter mechanism, ex vivo treatment of PDTFs with the bispecific antibodies failed to replicate the T cell activation observed with nivolumab in the same tumors, demonstrating that these molecules do not function as regular antagonists (**Fig. 1F**).

To directly investigate a potential PD-1-agonistic mechanism, anti-CD3/anti-CD28-stimulated PBMCs from healthy donors were cultured in the presence of increasing concentrations of αPD-1 (nivolumab), αCD4xαPD1, or αCD8xαPD1, respectively. In line with the fact that no substantial PD-L1 expression is present on PBMCs to inhibit T cell activity, nivolumab had no impact on T cell activation marker expression (CD137 on CD8^+^ T cells, OX40 on CD4^+^ T cells), T cell degranulation (CD107a), or T cell cytokine production (IFN-γ) (**Fig. 1, G to I**). In contrast, αCD4xαPD1 selectively suppressed the activation, degranulation, and cytokine production of CD4^+^ T cells without affecting CD8^+^ T cells, whereas αCD8xαPD1 inhibited these functions in CD8^+^ T cells, while leaving CD4^+^ T cells unaffected. Of note, αCD4xαPD1 and αCD8xαPD1 treatment of PDTFs did not reduce the expression of the activation markers OX40 or CD137 below baseline (**Fig. 1F**). Conceivably, due to the presence of PD-1 ligands in tumor tissues, PD-1 signaling does already take place and inclusion of the PD-1 agonist antibodies therefore does not further suppress T cell activity. Together, these findings indicate that αCD4xαPD1 and αCD8xαPD1 bispecific antibodies act as selective PD-1 agonists, leading to PD-1 signaling in CD4^+^ and CD8^+^ T cell subsets.

### CD8^+^ and CD4^+^ T cells independently initiate TME remodeling upon PD-1 blockade

While αCD8xαPD1 and αCD4xαPD1 do not functionally interrupt PD-1 signaling, they potently block PD-1 receptor accessibility on the cell surface. To selectively direct nivolumab to intratumoral CD4^+^ or CD8^+^ T cells, PDTFs were therefore preincubated with αCD8xαPD1 or αCD4xαPD1 to mask PD-1 on one subset, followed by nivolumab administration to inhibit PD-1 signaling on the opposing subset (**Fig. 2A**). This strategy was applied to a cohort of 21 tumors of various tumor types, including melanoma, ovarian cell carcinoma, non-small cell lung cancer, breast cancer and renal cell carcinoma, of which responsiveness to ex vivo PD-1 blockade had previously been established. In line with expectations, ex vivo nivolumab treatment resulted in increased activation of both intratumoral CD4^+^ and CD8^+^ T cells (**fig. S1A**). Moreover, pretreatment with both bispecifics (dual inhibition) confirmed that PD-1 receptor masking prevented the activation of both T cell subtypes as well as the initiation of a downstream cytokine and chemokine response (**fig. S1, A and B**). Directing PD-1 blockade to CD4^+^ T cells (αCD8xαPD1 + nivolumab; αPD-1➔CD4) resulted in a selective increase of the T cell activation marker OX40 on CD4^+^ T cells. In contrast, PD-1 blockade direction to CD8^+^ T cells (αCD4xαPD1 + nivolumab; αPD-1➔CD8) led to a selective increase of CD137 on CD8^+^ T cells (**Fig. 2, B and C**). These findings demonstrate that the bispecific approach developed here effectively isolates αPD-1 responses to individual T cell subsets.

**Figure 2.**
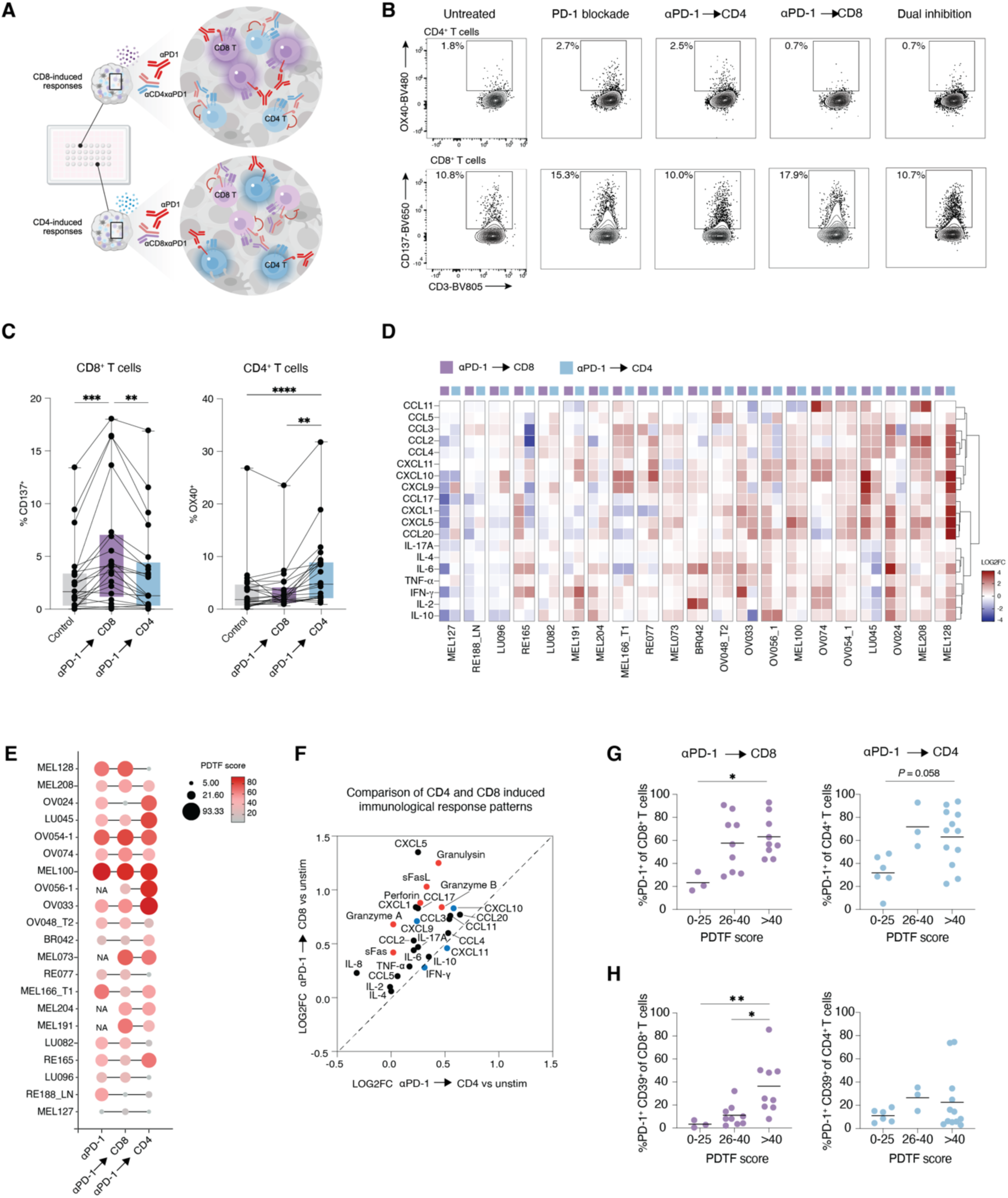
Capacity of both CD8^+^ and CD4^+^ T cells to induce TME remodeling following PD-1 blockade. (**A**) Schematic overview of experimental strategy. To direct PD-1 blockade specifically to CD8^+^ T cells (αPD-1➔CD8), PDTFs were pretreated with αCD4xαPD1 for 2 hours to block the PD-1 receptor on CD4^+^ T cells before administration of αPD-1 for 46 additional hours. The same strategy, but then using the αCD8xαPD1 bispecific antibody, was used to direct PD-1 blockade specifically to CD4^+^ T cells (αPD-1➔CD4). (**B**) Representative flow cytometry plots (MEL166-1) of OX40 expression in CD4^+^ T cells and CD137 expression in CD8^+^ T cells from PDTFs treated with αPD-1 (nivolumab), αPD-1➔CD4, αPD-1➔CD8, and both bispecific antibodies combined with αPD-1 (dual inhibition). (**C**) Quantification of CD137 on CD8^+^ T cells (left) and OX40 on CD4^+^ T cells (right) in PDTFs from αPD-1➔CD8 and αPD-1➔CD4 treated PDTFs, as measured by flow cytometry (n = 19). Treated conditions were compared to the dual inhibition condition (αPD-1 + αCD8xαPD1 + αCD4xαPD1), and in one case (OV054-1) to the unstimulated condition, where the dual inhibition condition did not pass quality control. \*\*\*\**P* < 0.0001, \*\*\**P* < 0.001, \*\**P* < 0.01 by Friedman test corrected for multiple comparisons. (**D**) Heatmap displaying log2 fold change (LOG2FC) values (αPD- 1➔CD4 and αPD-1➔CD8 conditions compared to the dual inhibition condition) of 19 soluble mediators secreted by PDTFs (n = 21). Parameters were ordered according to unsupervised hierarchical clustering. (**E**) Bubble plot representing the PDTF score developed by Voabil et al. (*18*), based on the most discriminative parameters between ex vivo αPD-1 responders and non- responders, consisting of IFN-γ, IL-10, CXCL9, CXCL10, CXCL11, CXCL5, CCL17, CCL4, CCL5, CCL20 and CXCL1 (n = 21). NA is displayed for samples that did not pass quality control. (**F**) Correlation of the averaged LOG2FC of soluble mediators secreted by PDTFs treated with αPD-1➔CD4 versus αPD-1➔CD8. Cytotoxic markers are indicated in red, markers related to the IFN-γ pathway in blue. (**G**) Percentage of PD-1^+^ cells and (**H**) PD- 1^+^CD39^+^ double positive cells in CD8^+^ and CD4^+^ T cells for each tumor, shown separately for tumors with a low (0–25), intermediate (26–40) or high PDTF score (>40) in the αPD-1➔CD8 condition and in the αPD-1➔CD4 condition.

To elucidate the contributions of intratumoral CD4^+^ and CD8^+^ T cells to the broad remodeling of the TME that occurs upon PD-1 blockade (*18*), this system was leveraged to characterize cytokine and chemokine profiles elicited by αPD-1-induced reinvigoration of CD8^+^ versus CD4^+^ T cells (**Fig. 2D**). Intriguingly, we observed that CD4^+^ and CD8^+^ T cells could both individually induce these downstream immunological responses. To better understand the individual impact of CD8^+^ and CD4^+^ T cells, we calculated PDTF response scores for each treatment condition. This score derives from changes in the parameters that are most discriminative between αPD-1 responding and non-responding tumors and was previously found to strongly correlate with clinical outcome (*18*). Of note, within tumors that are responsive to αPD-1 (nivolumab), we observed substantial variation in PDTF scores following selective CD8^+^ and CD4^+^ T cell reinvigoration, suggesting that in some tumors both subsets contribute similarly, whereas in other tumors, the TME response was predominantly driven by either CD8^+^ or CD4^+^ T cells (**Fig. 2E**). Log2-fold change (LOG2FC) analysis of soluble mediator profiles induced by αPD-1➔CD4 versus αPD-1➔CD8 revealed similar cytokine and chemokine patterns induced by reactivation of either T cell lineage but—in line with expectations—a higher secretion of cytotoxic mediators upon reactivation of CD8^+^ T cells (**Fig. 2F**).

To understand whether the different T cell reactivation patterns across tumors correlated with distinct abundance of specific states, we next compared the fraction of PD-1 expressiong CD8^+^ and CD4^+^ T cells. To this end, tumors were divided based on either high (>40), intermediate (26–40), or low (0–25) PDTF scores, respectively, reflecting either strong, weak or no TME remodeling. While the fraction of PD-1^+^ cells was low for both subsets in tumors with low PDTF scores, similarly high fractions were observed between tumors with intermediate and high scores (**Fig. 2G**). As further heterogeneity exists within the PD-1^+^ subset, we next assessed the fraction of cells co-expressing PD-1 and CD39 as a proxy for potentially tumor-reactive and dysfunctional cells (*46, 55*) (**fig. S1C)**. This revealed a clear trend for CD8^+^ T cells, with increasing fractions of PD-1^+^CD39^+^ cells in tumors with intermediate and high PDTF score, respectively (**Fig. 2H**). In contrast, fractions of double-positive cells were generally lower and more variable for CD4^+^ T cells. Altogether, these data suggest that upon reinvigoration by PD- 1 blockade both CD8^+^ and CD4^+^ T cells can independently induce broad TME remodeling in responding tumors, and that this response correlates with the increased presence of PD-1^+^ T cell populations.

### Targets of anti-PD-1 at the tumor site exhibit a highly dysfunctional state

To further characterize the differentiation state(s) of intratumoral PD-1^+^ T cells as potential targets of anti-PD-1 therapy, we performed single cell RNA and TCR sequencing (scRNA+TCR-seq) on T cells from eight tumors. For both T cell lineages, clustering analysis of the CD8^+^ and CD4^+^ T cell compartment revealed one cluster with high *PDCD1* gene expression (encoding PD-1) that was present across all tumor samples (**Fig. 3A and fig. S2A**). Both *PDCD1^+^* clusters co-expressed *CXCL13* and *ENTPD1* (encoding CD39) as markers for tumor reactivity (*41, 46, 56, 57*), although expression of the latter was lower in the CD4^+^ cluster (**Fig. 3A**). The CD4^+^ subset also contained a second *ENTPD1^+^* cluster related to *FOXP3^+^* regulatory T cells (Tregs) that showed low *PDCD1* expression (**Fig. 3A, fig. S2B**), as reported previously (*58, 59*). Analysis of differentially expressed genes (DEG) further revealed that *PDCD1*^+^ CD8^+^ and CD4^+^ T cells shared a core dysfunctional program (**Fig. 3, B and C**), in line with prior observations (*59, 60*). This program included upregulation of *TOX*, *HAVCR2* (encoding TIM-3), *LAG3*, *CXCL13*, *TNFRSF9* (CD137) or *TNFRSF4* (OX40), and *IFNG* (**Fig. 3D and Table S3**), consistent with hallmarks of T cell exhaustion reported in LCMV models (*6*). Of note, the enrichment of *CXCL13* in both subsets highlights the unique aspects of T cell exhaustion in human tumor tissues (*56, 61*). Moreover, *PDCD1*^+^ CD8^+^ and CD4^+^ T cells exhibited gene signatures not only associated with exhausted but also with effector T cells (*33, 62*), while showing reduced scores for naïve (*62*) and progenitor gene signatures (*63*) compared to their *PDCD1*^−^ counterparts (**Fig. 3E, fig. S2C**). In line with this subset containing tumor-reactive T cells, *PDCD1^+^*cells displayed higher expression of signatures for TCR signaling (Reactome) and neoantigen specificity (*30*), as well as increased TCR clonality (**fig. S2, C and D**).

**Figure 3.**
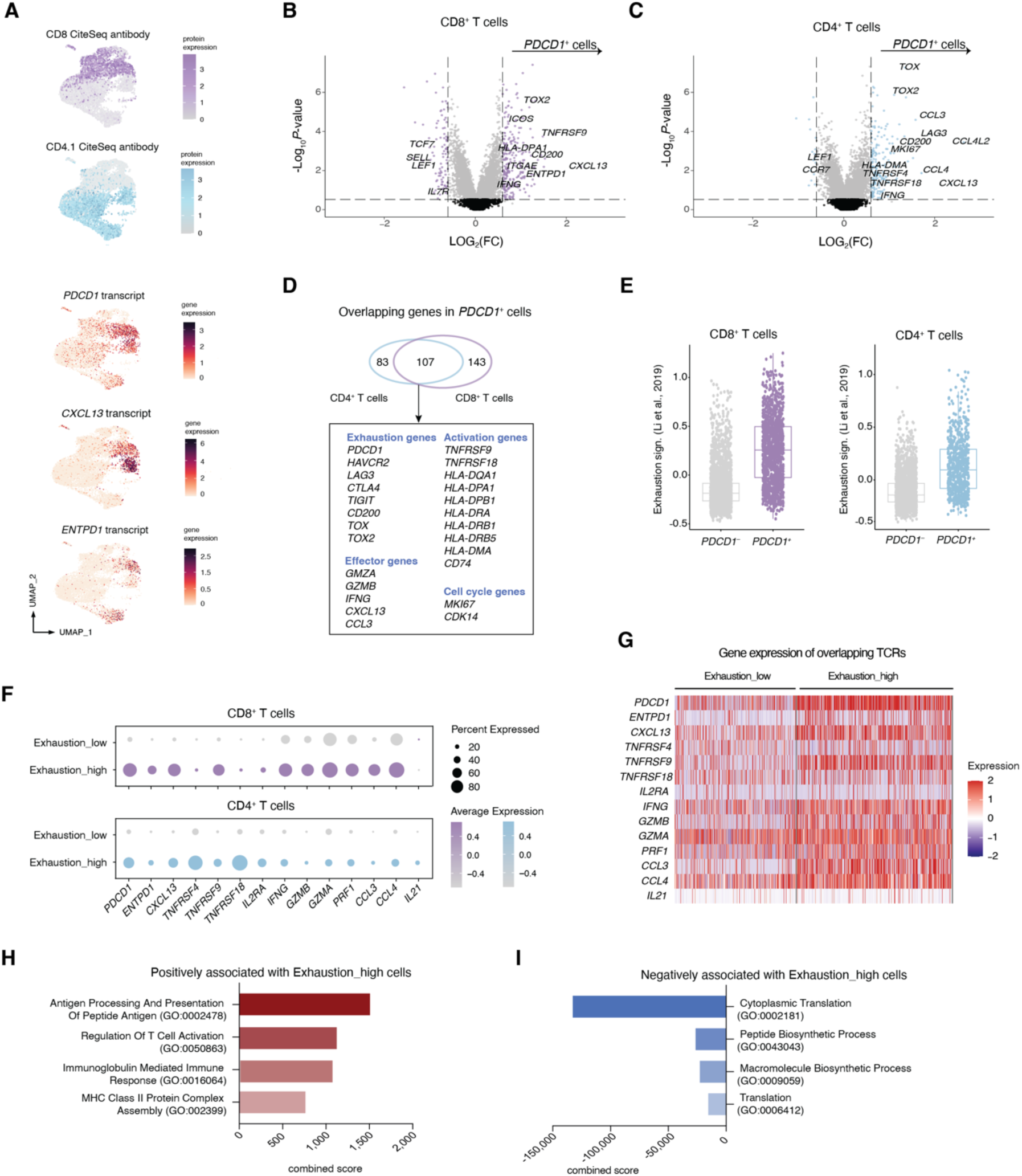
Characterization of intratumoral PD-1 expressing CD8^+^ and CD4^+^ T cells as potential targets of αPD-1. (**A**) UMAPs of intratumoral T cells displaying expression of CD8 and CD4.1 measured by CITE-seq antibody binding, and *PDCD1* (encoding PD-1), *CXCL13* and *ENTPD1* (encoding CD39) transcript expression (n = 8). (**B+C**) Differential gene expression analysis (pseudo-bulk, n = 8) between *PDCD1^+^* and *PDCD1*^−^ CD8^+^ (**B**) and CD4^+^ T cells (**C**). (**D**) Number of individual and overlapping genes between *PDCD1^+^* intratumoral CD4^+^ and CD8^+^ T cells, with overlapping key genes indicated. (**E**) Expression of an exhaustion gene signature (*33*) in *PDCD1^+^* and *PDCD1^−^*intratumoral CD8^+^ and CD4^+^ T cells. (**F**) Expression of T cell effector genes in CD8^+^ and CD4^+^ T cells separated into Exhaustion_high (top 10% cells) and Exhaustion_low (remaining 90% cells) groups, according to the gene signature in (**E**) adjusted by removal of four effector genes *PDCD1*, *IFNG*, *TNFRSF9* and *CCL3*. **(G)** Heatmap showing a similar analysis as in **(F)** but including only T cells with TCRs that overlap between Exhaustion_low and Exhaustion_high subgroups. (**H+I**) Gene Ontology (GO) enrichment analysis of pathways positively (**H**) or negatively (**I**) associated with Exhaustion_high as compared to Exhaustion_low T cells. X-axis displays a combined score calculated as log(*P*-value) x z-score.

During their differentiation from precursors, dysfunctional T cells gradually lose effector functions (*64*). We, however, detected increased effector gene expression in the *PDCD1^+^*subsets compared to their *PDCD1^−^* counterparts, a feature previously observed in exhausted T cells in both chronic infections and tumors (*6, 32, 33, 64, 65*). As suggested by the presence of both CD39^+^ and CD39^−^ cells within the PD-1^+^ population (**Fig. 2, G and H**), *PDCD1^+^* T cells may comprise multiple cell states with different levels of dysfunction. Therefore, to examine whether effector potential, as measured by mRNA abundance, is decreased in late- dysfunctional cells, we compared highly exhausted (top 10% exhaustion signature (*33*)) to the remaining T cells. Notably, transcripts for T cell effector, activation, and cytotoxicity genes, such as *TNFRSF9*, *IFNG*, *GZMA*, *GZMB*, and *PRF1,* were all significantly more abundant in the highly exhausted cells (**Fig. 3F, fig. S2E**). As the less exhausted population can include a substantial proportion of bystander cells, we also compared effector gene expression in exhaustion_low and exhaustion_high cells that share TCR sequences. This analysis similarly revealed higher effector gene expression in the exhaustion_high subset compared to their exhaustion_low counterparts (**Fig. 3G**). To further investigate this disconnect between gene expression and function, we performed gene ontology (GO) enrichment analysis. This confirmed that highly exhausted T cells were substantially higher in T cell activation and immune response pathways (**Fig. 3H**). Strikingly, exhaustion_high T cells did score significantly lower for pathways related to cytoplasmic translation and peptide biosynthesis than their less exhausted counterparts (**Fig. 3I**). This pattern was also observed when the same analysis was restricted to cells with shared TCRs (**fig. S2, F and G**). These results suggest that dysfunctional T cells display a highly activated transcriptional state in tumor tissues, likely driven by continuous antigen stimulation, but may have reduced translational activity, potentially preventing effective protein synthesis and ultimately effector function.

### Intratumoral T cells do not undergo transcriptional state changes in response to PD-1 blockade

Having identified T cells with elevated PD-1 expression and characteristics of both effector capacity and dysfunction as potential primary targets of PD-1 blockade at the tumor site, we next sought to understand whether and how this T cell pool is reinvigorated by αPD-1 treatment. To this end, PDTFs from six αPD-1-responsive tumors were treated ex vivo for 48 hours, followed by scRNA+TCRseq of sorted CD45^+^ immune cells (**Fig. 4A**). In line with prior data, PD-1 blockade increased secreted IFN-γ and its downstream mediators CXCL9, CXCL10, and CXCL11 (**Fig. 4B**). Furthermore, pretreatment with an Lck inhibitor (Lcki), which blocks downstream TCR-signaling, prevented secretion of all mediators, verifying that intratumoral T cells are driving the observed TME remodeling. Additionally, pretreatment with an anti-IFN-γ receptor antibody (αIFNγR) effectively reduced the levels of IFN downstream chemokines without affecting IFN-γ secretion itself, further validating the functionality of the system.

**Figure 4.**
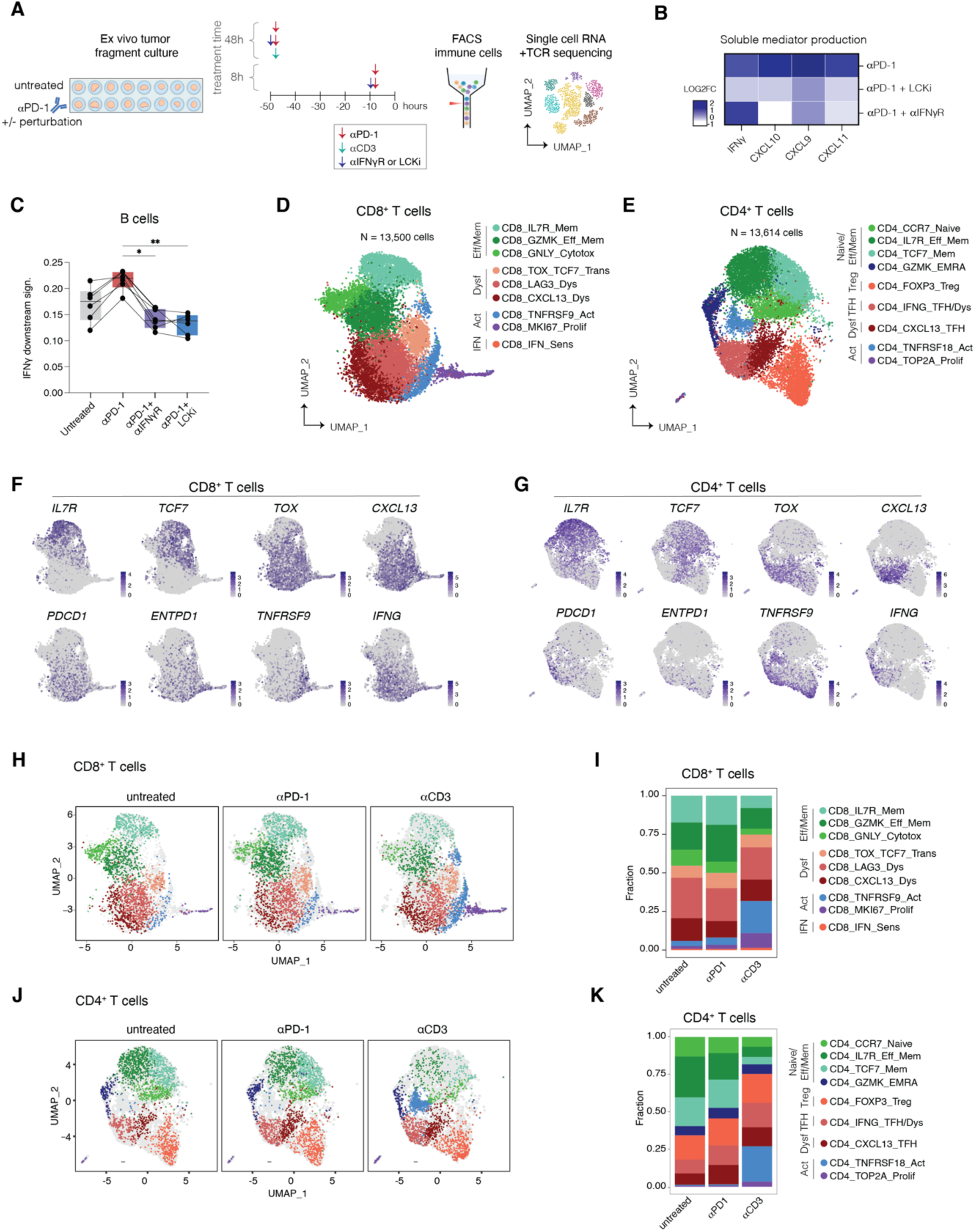
Lack of major cell state remodeling of intratumoral T cells in response to anti- PD-1. (**A**) Schematic overview of the experimental setup using the PDTF platform combined with flow sorting of CD45^+^ immune cells and subsequent single-cell RNA sequencing. (**B**) Heatmap displaying the mean LOG2FC (treated condition divided by the untreated condition for each tumor) of secreted IFN-γ and downstream mediators CXCL9, CXCL10 and CXCL11 from PDTFs treated with αPD-1, αPD-1+αIFNγR, and αPD-1+Lck inhibitor (Lcki) versus untreated PDTFs. (**C**) IFN-γ response gene signature (*33*) in B cells separated per tumor and experimental condition (n = 6). \*\**P* < 0.01, \**P* < 0.05 by Friedman test corrected for multiple comparisons. (**D**) UMAP visualization of all intratumoral CD8^+^ T cells (from untreated and all treated conditions) in human tumor fragments from six tumors (n = 13,500 cells) identifying nine different clusters. (**E**) UMAP visualization of all intratumoral CD4^+^ T cells (from untreated and all treated conditions) in human tumor fragments from six tumors (n = 13,614 cells) identifying nine different clusters. (**F**) UMAPs of intratumoral CD8^+^ T cells and (**G**) of CD4^+^ T cells displaying expression of different canonical marker genes (n = 6 tumors). (**H**) UMAPs displaying CD8^+^ T cells from untreated, αPD-1-treated, and αCD3-treated (both 48 hours) conditions, down-sampled to equal cell numbers per condition. (**I**) Bar graphs of cluster fractions of CD8^+^ T cell states derived from untreated, αPD-1-treated and αCD3-treated conditions. (**J**) UMAPs displaying CD4^+^ T cells from untreated, αPD-1-treated and αCD3- treated conditions, down-sampled to equal cell numbers per condition. (**K**) Bar graphs of cluster fractions of CD4^+^ T cell states derived from untreated, αPD-1-treated and αCD3-treated conditions.

To assess whether the observed TME remodeling could also be captured at the transcriptional level, B cells isolated at different timepoints post ex vivo anti-PD-1 therapy were analyzed by scRNAseq. Intratumoral B cells do not form direct targets of αPD-1 as they largely lack PD-1 expression (**fig. S3A**), but are capable of sensing secreted cytokines and responding to these

(*66*). Notably, 48 hrs after treatment, B cells showed upregulation of a gene signature associated with IFN-γ sensing (*33*) (**Fig. 4C**), while this signature was reduced below baseline levels following pretreatment with αIFNγR and Lcki, respectively. Interestingly, even a brief 8 hr treatment with αPD-1 (resulting in complete PD-1 engagement, **fig. S3B**), already induced detectable IFN-γ sensing in B cells, indicating that intratumoral T cells can initiate a tissue response within hours of αPD-1 administration (**fig. S3C**).

Having established that PD-1 blockade induced a rapid sensing of IFN-γ by the B cell compartment, we next aimed to characterize the treatment-induced changes in the T cell compartment. Therefore, we first identified distinct CD8^+^ and CD4^+^ phenotypes by separate clustering analysis of each subset. Both CD8^+^ and CD4^+^ T cells each clustered in nine distinct transcriptional states (**Fig. 4D to G**), including naïve/memory-like, effector, dysfunctional, proliferating and activated clusters. A follicular helper T cell (TFH) and a regulatory T cell (Treg) cluster were exclusive to CD4^+^ T cells (**fig. S4, A to D**). To link this dataset to our prior analysis, we plotted the gene signatures of *PDCD1⁺*CD4⁺ and CD8⁺ T cells (**Fig. 3D**) on the cultured PDTF dataset. The most prominent signature expression was observed in the dysfunctional and activated clusters (**fig. S5A**). Further characterization of the dysfunctional T cell compartment identified one early-dysfunctional/transitional (CD8_TOX_TCF7_Trans) and two late-dysfunctional (CD8_LAG3_Dys, CD8_CXCL13_Dys) clusters in CD8^+^ T cells, as well as one late-dysfunctional CD4^+^ cluster (CD4_IFNG_TFH/Dys), based on marker gene expression and cross-labeling with a terminal exhaustion signature (*33*) (**fig. S5, B to E**). All late-dysfunctional states also showed elevated expression of a tumor reactivity signature (*29*) and increased clonal expansion (**fig. S5, F to H**), in line with potential tumor recognition. Of note, we did not observe precursor or intermediate dysfunctional states intratumorally which have previously been associated with ICB response in chronic infection mouse models (**fig. S5C and D**) (*39, 67*). Consistent with our first dataset, both late-differentiated CD8^+^ and CD4^+^ cell states showed high expression of activation/ effector genes, such as *TNFRSF9* and *IFNG* (**Fig. 4, F and G**), suggesting that they acquire both an activated and highly dysfunctional phenotype.

We next aimed to characterize the treatment-induced changes in these T cell states. To this end, alterations in cell-state fractions were quantified upon 48-hour treatment with αPD-1. This analysis revealed that αPD-1 treatment elicited minimal cluster changes within the CD8^+^ T cell compartment compared to untreated fragments (**Fig. 4, H and I and fig. S6A**). αCD3 was used as a positive control to benchmark the changes induced by TCR triggering in the entire tumor-resident T cell compartment, regardless of differentiation state or tumor reactivity. In contrast to αPD-1, αCD3 treatment resulted in substantial remodeling of the intratumoral T cell landscape, prominently inducing the CD8_TNFRSF9_Act and CD8_MKI67_Prolif clusters (**Fig. 4I and fig. S6A**). A similar lack of consistent cell state shifts upon αPD-1 treatment was observed in CD4^+^ T cells, while αCD3 stimulation also led to the emergence of an activation-specific cluster CD4_TNFRSF18_Act and, to a lesser extent, a proliferating cluster CD4_TOP2A_Prolif (**Fig. 4, J and K and fig. S6B**). Notably, αCD3-induced activation coincided with a significant decrease in memory and effector-memory clusters (CD8_IL7R_Mem, CD8_GZMK_Eff_Mem, CD8_GNLY_Cytotox for CD8^+^ T cells; CD4_TCF7_Mem and CD4_IL7R_Eff_Mem for CD4^+^ T cells), while the more dysfunctional clusters did not substantially decrease (**fig. S6C**). This suggests that dysfunctional T cell states may be less responsive to transcriptional reprogramming upon TCR stimulation, in line with their fixed epigenetic state (*50, 68, 69*).

### Tumor-residing T cells do not show transcriptional rewiring but are activated at the protein level by αPD-1

To identify more nuanced changes induced by αPD1 or αCD3 treatment, we performed DEG analysis. This analysis revealed no significantly differentially expressed genes between untreated and αPD-1-treated PDTFs on CD8^+^ T cells, whereas αCD3 treatment led to significant upregulation of numerous activation markers, including *IL2RA*, *GZMB*, and *LAG3* (**Fig. 5, A and B**). The same pattern was observed in the CD4^+^ compartment, identifying limited transcriptional alterations in response to αPD-1, with only *LTA*, *IL2*, and *IL21* emerging as differentially expressed genes (**Fig. 5, C and D, fig. S6D**). Additionally, DEG analysis after 8 hours of αPD-1 treatment for both T cell compartments confirmed these findings, indicating no significant transcriptional alterations (**fig. S6E)**.

**Figure 5.**
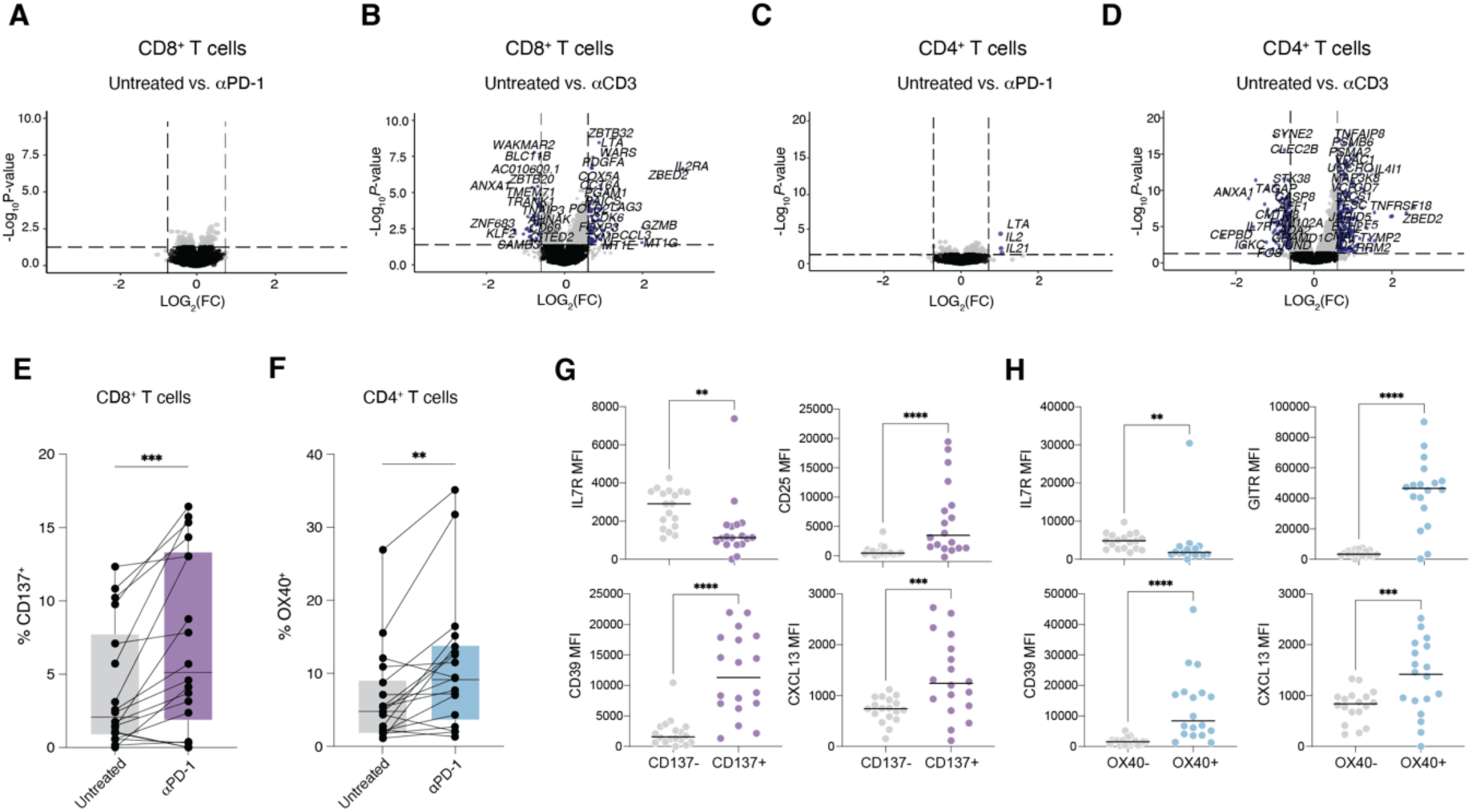
Dysfunctional intratumoral T cells display changes in response to PD-1 blockade predominantly at the protein level. (**A-B**) Differential gene expression analysis in CD8^+^ T cells (pseudo-bulk, n = 6) between the untreated and αPD1-treated (**A**) or αCD3-treated (**B**) condition. (**C-D**) Differential gene expression analysis in CD4^+^ T cells (pseudo-bulk, n = 6) between the untreated and αPD1-treated (**C**) or αCD3-treated (**D**) condition (48 hours). (**E**) Quantification of CD137 on total intratumoral CD8^+^ T cells in untreated and αPD-1-treated PDTFs, as measured by flow cytometry (n = 18). \*\*\**P* < 0.001 by Friedman test corrected for multiple comparisons. (**F**) Quantification of OX40 on total intratumoral CD4^+^ T cells in untreated and αPD1-treated PDTFs, as measured by flow cytometry (n = 18). \*\**P* < 0.01 by Friedman test corrected for multiple comparisons. (**G**) Difference in IL7R, CD39, CD25 and CXCL13 expression, as shown by mean fluorescence intensity (MFI), in CD137^−^ and CD137^+^ CD8^+^ T cells from αPD-1-treated PDTFs (n = 18). \*\**P* < 0.01, \*\*\**P* < 0.001 and \*\*\*\**P* < 0.0001 by Wilcoxon matched-pairs test. (**H**) Difference in IL7R, CD39, GITR and CXCL13 expression, as shown by mean fluorescence intensity (MFI), in OX40^−^ and OX40^+^ CD4^+^ T cells from αPD-1-treated PDTFs (n = 18). \*\**P* < 0.01, \*\*\**P* < 0.001 and \*\*\*\**P* < 0.0001 by Wilcoxon matched-pairs test.

In contrast to the limited transcriptional changes following PD-1 blockade, αPD-1 treatment did result in a significant increase in CD137 and OX40 protein expression on both CD8^+^ and CD4^+^ T cells (**Fig. 1, A and B and Fig. 5, E and F**). Furthermore, comparison of CD137^+^ and CD137^−^ cells from αPD-1-treated PDTFs demonstrated that the reinvigorated CD8^+^ T cells expressed markers characteristic of activated and dysfunctional T cells, including CD25, CD39, CXCL13, GITR, as well as reduced IL7R expression (**Fig. 5G and fig. S7A**). Similarly, reinvigorated CD4^+^ T cells displayed increased levels of markers associated with T cell dysfunction, including CD39 and CXCL13, alongside decreased IL7R expression (**Fig. 5H and fig. S7B**). Additionally, these reactivated cells also expressed Treg-associated markers, such as FOXP3, CD25, and GITR.

Collectively, these data demonstrate that strong TCR engagement via αCD3 can lead to transcriptional remodeling of the intratumoral T cell landscape, which appears mostly driven by activation of non-dysfunctional (likely bystander) T cells. In contrast, αPD-1 treatment induces only minimal transcriptional changes in both intratumoral CD8^+^ and CD4^+^ T cells, despite evidence of a downstream functional immune response. Importantly, T cell activation following αPD-1 treatment is clearly detectable at the protein level within the dysfunctional compartment, suggesting that PD-1 blockade may exert its effect post-transcriptionally.

### PD-1 blockade increases translation capacity in dysfunctional T cells

Considering that αPD-1-induces T cell-dependent remodeling of the TME in the absence of appreciable transcriptional changes, we hypothesized that T cell reactivation at the tumor site may primarily be driven at the post-transcriptional level rather than through transcriptional rewiring. In line with this idea, both CD8^+^ and CD4^+^ dysfunctional T cells in tumors showed the highest mRNA expression of effector genes like *IFNG* (**Fig. 4F and G**). Notably, comparing IFN-γ protein levels in the supernatant of PDTFs with corresponding *IFNG* transcript levels in T cells from the same tumors, we observed that PD-1 blockade led to a marked increase in IFN- γ protein without a concomitant rise in *IFNG* mRNA expression (**Fig. 6A**).

**Figure 6.**
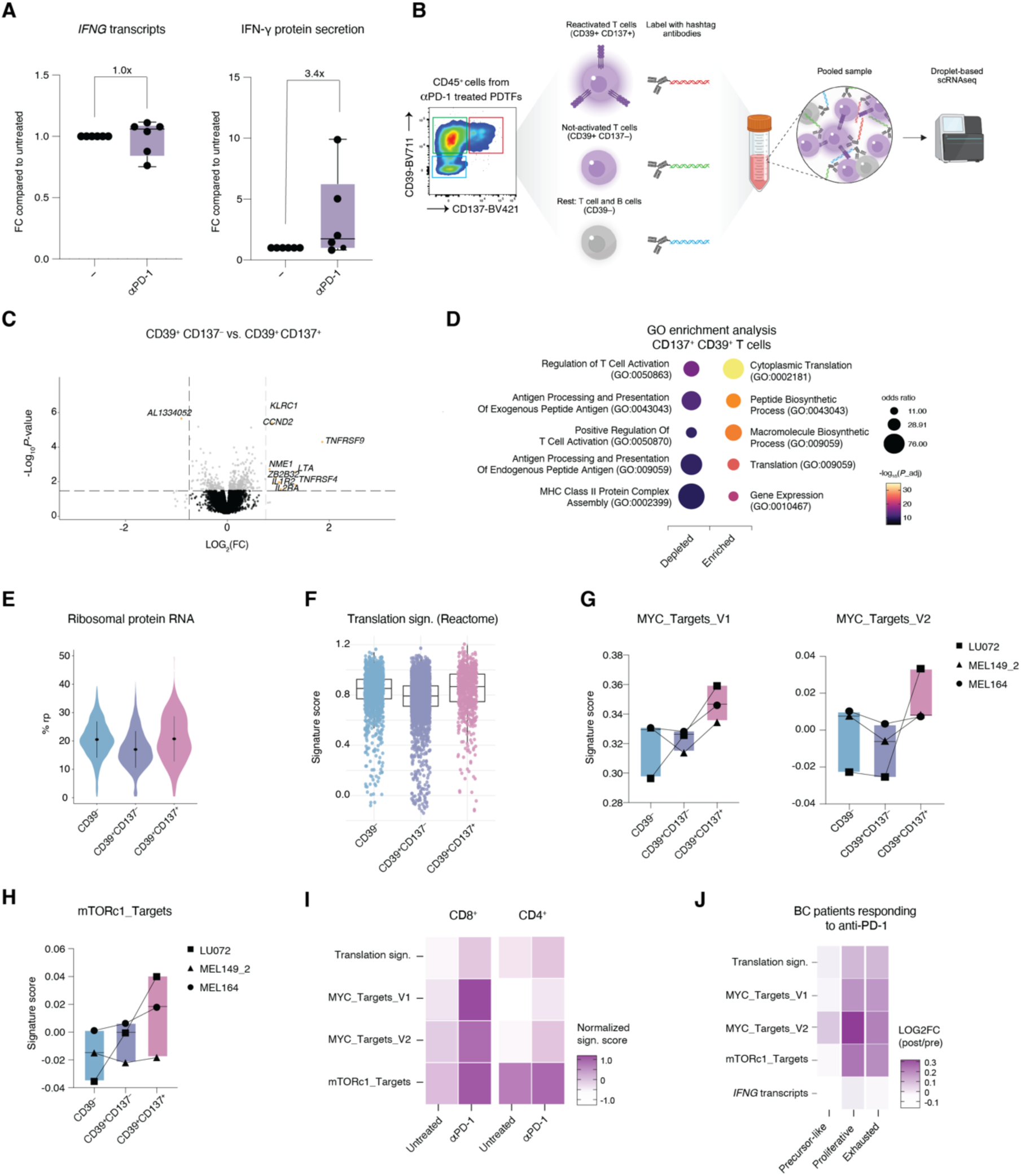
PD-1 blockade restores translational capacity in intratumoral dysfunctional T cells. (**A**) *IFNG* gene expression in total T cells (left) and IFN-γ protein secretion measured in the supernatants (right) from the same PDTFs that were left untreated and treated with αPD-1, with the fold change displayed on top. (**B**) Schematic overview of the experimental design to isolate reactivated dysfunctional (CD39^+^CD137^+^), non-reactivated dysfunctional (CD39^+^CD137^−^) T cells and remaining T and B cells (CD39^−^). To capture the earliest changes and enrich for higher cell numbers, a shorter incubation time of 12 hours was used. (**C**) Differential gene expression analysis (pseudo-bulk, n = 3) between non-reactivated CD39*^+^*CD137^−^ and reactivated CD39^+^ CD137^+^ T cells. (**D**) Gene ontology enrichment analysis comparing reactivated CD39^+^ CD137^+^ T cells to non-reactivated CD39^+^ CD137^−^ T cells. (**E**) Percentage of ribosomal protein RNA as a measurement of translation capacity in the CD39^+^ CD137^+^ activated, CD39^+^ CD137^−^, and CD39^−^ T cell populations. (**F**) Expression of a translation signature (Reactome) in the CD39^+^ CD137^+^ activated, CD39^+^ CD137^−^, and CD39^−^ T cell populations. (**G**) MYC_Targets_V1, MYC_Targets_V2, and (**H**) mTORc1_Targets (Reactome) gene signature scores in CD39^+^CD137^+^, CD39^+^CD137^−^, and CD39^−^ T cell populations per tumor. (**I**) Heatmap depicting normalized gene signature scores for Translation, MYC_Targets_V1, MYC_Targets_V2, and mTORc1_Targets (Reactome) in intratumoral *ENTPD1^+^*(CD39) CD8⁺ T cells (left) and CD4⁺ T cells (right) following 8 hr ex vivo PD-1 blockade. (**J**) Heatmap of LOG2FC of the gene signature scores for Translation, MYC_Targets_V1, MYC_Targets_V2, mTORc1_Targets (Reactome) and *IFNG* transcripts in intratumoral CD8^+^ T cell clusters (assigned by Liu et al., 2022 (*56*)) from breast cancer patients treated with anti-PD-1 in post- versus pre-treatment samples (*74*).

Next, to directly assess changes within the reinvigorated T cell population, we sorted non-reactivated and reactivated T cells based on CD137 expression after ex vivo culture with PD- 1 blockade (n = 3), using CD39 as a stable marker of dysfunction that is unaffected by PD-1 blockade (**Fig. 6B and fig. S7, C and D**). Despite limited cell numbers (**fig. S7E**), DEG analysis could be performed and again revealed minimal transcriptional differences between reactivated dysfunctional (CD39^+^ CD137^+^) and non-reactivated dysfunctional (CD39^+^ CD137^−^) T cells from αPD-1 treated PDTFs (**Fig. 6C**). Strikingly, reactivated T cells were enriched in pathways related to cytoplasmic translation and biosynthesis processes, in line with the notion that activation in response to PD-1 blockade may be mediated primarily through an increase in translational capacity (**Fig. 6D**). Of note, non-activated CD39^+^ cells exhibited enrichment in T cell activation pathways, though this was primarily driven by the higher expression of HLA class II genes and showed lower statistical significance compared to the translation-related pathways in reactivated cells. This also highlights the challenge in distinguishing truly reactivated cells from dysfunctional CD39^+^ cells based solely on transcriptional profiles.

Consistent with the enrichment for cytoplasmic translation, reinvigorated CD137^+^ T cells showed a modest trend towards increased expression of a translation signature (Reactome) as well as in the percentage of ribosomal protein-encoding transcripts compared to their non-reactivated T cell counterparts, suggesting—at least partial—upregulation of the translational machinery upon reinvigoration (**Fig. 6, E and F**). Previous studies highlighted the relevance of MYC and mTOR signaling specifically for T cell effector protein translation capacity (*70–73*). Accordingly, reinvigorated dysfunctional T cells displayed a marked upregulation of gene signatures associated with MYC and mTORc1 signaling (**Fig. 6, G and H and fig. S7F**). To evaluate their relevance for T cell activation, these signatures were also assessed within PDTFs treated with αCD3. Supporting this notion, the αCD3-induced activated and proliferating clusters (CD8_TNFRSF9_Act, CD8_MKI67_Prolif, CD4_TRFRSF18_Act and CD4_TOP2A_prolif) exhibited the highest scores for these signatures (**fig. S7, G and H**). We further validated these findings using our dataset of PDTFs treated with αPD-1 for 8 hours, focusing on CD39 (encoded by *ENTPD1*)-positive T cell subsets (**Fig. 6I**). In CD8⁺ T cells, αPD-1 treatment induced an increase in all four translation-associated signatures within the late dysfunctional CD39⁺ subset. In CD4⁺ T cells, three of these four signatures exhibited a modest increase in the CD39^+^ subset.

Lastly, to understand the clinical relevance of these observations, we examined the translation-associated gene signatures in a clinical scRNAseq data set from breast cancer patients obtained pre- and post-anti-PD-1 treatment (*74*). Post-treatment samples demonstrated an increase in translation-associated gene signatures predominantly within proliferating and exhausted T cell clusters, with no similar changes observed in precursor-like cells (**Fig. 6J**). Interestingly, these translation-related changes occurred without a corresponding rise in *IFNG* transcript levels across clusters. Furthermore, this increase in translational capacity was notably absent in non-responding patients, highlighting the link between increased translational activity in dysfunctional T cells and clinical anti-PD-1 response (**fig. S7I**).

Collectively, our results indicate that both dysfunctional CD4^+^ and CD8^+^ T cells at the tumor site can be reinvigorated by anti-PD-1 treatment via enhancement of their translational capacity. These data point towards a model in which dysfunctional CD4^+^ or CD8^+^ T cells acquire a translational block upon chronic antigen stimulation, which may be alleviated by PD-1 blockade. This restoration of effector protein translation capacity likely enables the dysfunctional T cell pool to initiate the local anti-tumor immune response following αPD-1 treatment.

## DISCUSSION

Although significant progress has been made in understanding the mechanism of action of PD- 1 blocking therapies, the precise incipient effects of this treatment on T cells in tumor tissues have remained incompletely understood. Our data reveal that T cells at the tumor site can induce potent immunological changes in other cell types within a few hours after PD-1 blockade initiation. Notably, both CD8^+^ and CD4^+^ T cells are capable to independently instigate remodeling of the TME upon anti-PD-1. These findings do not preclude additional mechanisms of CD4^+^ T cell-mediated help, such as dendritic cell activation via CD40L (*75*) that may also contribute to ICB response, but rather highlight a shared role of CD4^+^ and CD8^+^ T cells at the tumor site. Importantly, the T cells responding to anti-PD-1 treatment exhibit a highly dysfunctional phenotype, and our data are consistent with a model in which their reinvigoration predominantly occurs through restoration of translational capacity rather than transcriptomic rewiring. This process, which enables the rapid production of key effector molecules such as IFN-γ, highlights a novel mechanism for dysfunctional T cell activation in tumor tissues.

Several studies in murine models of cancer have identified precursor-like T cells as essential drivers of durable anti-tumor immunity (*14, 15, 76, 77*). Strikingly, our ex vivo analyses in human cancer tissues reveal that dysfunctional T cells are the key initiators of anti-PD-1 induced immune responses at the tumor site. While the latter observation is in line with the previously described association of dysfunctional T cells in pre-treatment biopsies with clinical outcome to PD-1 blockade (*41, 45, 46*), it challenges the notion that these cells have no capacity for reinvigoration because of their terminal, epigenetically fixed state (*48–50*). Recent work from the Amit lab reported that anti-PD-1 therapy in mouse models predominantly accelerates T cell differentiation from precursor to dysfunctional states without significant transcriptional reinvigoration of the latter (*78*). Accordingly, we observed no substantial transcriptional changes in dysfunctional T cells upon PD-1 blockade in human tumors. Nevertheless, activation of these cells was clearly detectable at the protein level. From a transcriptional point of view, dysfunctional T cells are well-equipped for effector function as they highly express a range of effector genes including *IFNG*, *GZMA*, *GZMB* and *CXCL13* (*6, 41, 64, 77, 79*). The lack of antitumor function despite high effector RNA expression observed in dysfunctional T cells therefore suggests that these cells may exhibit a block at the level of post-transcriptional regulation. Further supporting the notion that PD-1 blockade may act via restoration of translation capacity, T cell activation and downstream IFN-γ responses were detectable as early as eight hours post-treatment, suggesting the existence of a swift protein-level change in dysfunctional T cells. Collectively, these results lead to a model in which PD-1 blockade not only acts via accelerating the proliferation and differentiation of stem-like T cells (*78, 80*), but also by restoring the effector capacity of T cells with established dysfunction.

The importance of post-transcriptional regulation of T cell effector function, and especially of cytokines such as IFN-γ, is increasingly recognized, and our findings contribute to this growing body of evidence. Specifically, consistent with our observations, it has been demonstrated that memory CD8^+^ T cells contain pre-formed cytokine-encoding mRNA, which is translationally repressed but upon T cell activation enables rapid cytokine production (*81*). OT-1 T cells transferred into a murine B16-OVA melanoma model were found to display impaired IFN-γ protein secretion despite high mRNA expression due to post-transcriptional regulatory mechanisms (*82*). Interestingly, PD-1 blockade could also increase IFN-γ production in this setting. This post-transcriptional regulation mechanism is not exclusive to CD8^+^ T cells, as tissue-resident CD4^+^ T cells have also been reported to harbor mRNAs encoding proinflammatory cytokines in the absence of corresponding protein production (*83*). Recent work by Van Der Byl et al. (*84*) provides further insight into this phenomenon, showing that tolerant CD8^+^ T cells, which have entered a state of unresponsiveness after antigen recognition, exhibit impaired protein translation (*84*). Importantly, restoration of protein translation in these cells required MYC expression, in line with MYC’s established role as a key regulator of protein translation in effector T cells (*71, 85*). Consistent with these findings, we observed increased MYC target gene signatures in dysfunctional T cells reinvigorated by PD-1 blockade. While MYC is involved in general translation enhancement, mTOR appears to have both general and specific effects on protein synthesis in T cells (*86*). A recent study revealed a novel mechanism by which mTOR regulates cytokine production through 3’UTR-mediated translation control, particularly involving AU-rich elements in cytokine mRNAs (*73*). Moreover, in line with our data, studies in mouse models of chronic infection have shown that mTOR is crucial for the activation of stem-like CD8^+^ T cells following anti-PD-1 therapy (*67*). The upregulated mTOR signaling we observed may suggest that reinvigorated dysfunctional T cells prioritize the translation of transcripts encoding proteins necessary for T cell effector functions, such as cytokines and other immune mediators, a question that will be important to address in future research. Notably, this uncoupling of TCR signaling from protein translation observed in dysfunctional T cells may serve as a protective mechanism to prevent uncontrolled tissue damage in the context of persistent antigen stimulation and inflammation.

Our study has a number of limitations: First, the bispecific anti-PD-1 antibodies used did not allow for direct selective activation of CD4⁺ or CD8⁺ T cells, making it necessary to use an indirect approach, by agonizing PD-1 receptors on one of the T cell subsets of interest while blocking PD-1 engagement on remaining PD-1⁺ cells. As a result, binding of anti-PD-1 to other (non-T cell) populations cannot formally be excluded. However, observations from the dual inhibition experiments (**fig. S1**) suggest that such populations are unlikely to play a relevant functional role in initiating downstream immunological responses. Second, while our data support a functional role for CD4⁺ T cells in driving intratumoral anti–PD-1 responses, the extent to which CD4-mediated immunity contributes to clinical outcome remains to be determined. Finally, the PDTF model exclusively captures immune responses at the tumor site. However, compelling evidence has accumulated indicating that lymph node-resident precursor-like T cells are critical for sustaining anti–PD-1 responses (*14–17*), suggesting that key elements of the immune response take place outside the TME. Building on these previous studies and our findings, we propose a model in which dysfunctional and precursor-like T cells have an independent and synergistic role in driving the response to PD-1 blockade. Dysfunctional T cells by themselves are likely unable to mediate durable responses to ICB, because of the previously observed inability to undergo epigenetic reprogramming. Due to their short lifespan (*87*), constant replenishment by less dysfunctional T cells may be required to maintain the dysfunctional compartment at the tumor site. Compelling evidence from mouse models has shown that this replenishment is mediated by the stem-like progenitor T cells that mainly reside in lymphoid tissues (*14, 60*). Importantly, ICB both promotes a proliferative burst of progenitor T cells essential to drive long-lasting responses (*77, 88*) and accelerates their transition to more differentiated cell states in the TME (*78*). Our results reveal additional critical steps in the anti-PD-1 response driven by the reinvigoration of dysfunctional T cells in tumor tissues: First, the alleviation of the translational block in dysfunctional T cells may be critical for these cells to respond to PD-1 blockade, but also for ensuring effector function of replenished cells. Second, the reactivation of dysfunctional T cells at the tissue site may be essential for initiating cytokine and chemokine secretion, which fosters the continuous recruitment of progenitor T cells needed for durable antitumor immunity.

In conclusion, our study highlights tumor-residing dysfunctional CD8^+^ and CD4^+^ T cells as critical drivers of TME remodeling upon PD-1 blockade and unravels a new mechanism of their therapeutic reinvigoration via restoration of translational activity. These findings provide evidence that these cells are not merely ‘exhausted’ non-functional entities in the tumor landscape, but equipped to rapidly regain anti-tumor effector function in response to PD-1 blockade.

## MATERIALS AND METHODS

### Collection of patient-derived tumor fragments

Human tumor tissue was collected from cancer patients undergoing surgery for non-small cell lung cancer (LU), ovarian cancer (OV), melanoma (MEL), breast cancer (BR), and renal cell carcinoma (RE) at the Netherlands Cancer Institute/Antoni van Leeuwenhoek Hospital (NKI-AVL) between March 2020 and September 2024 (**Table S1**). The study (CFMPB484) was approved by the institutional review board of the NKI-AVL and performed in compliance with all ethical regulations. Patients provided prior written consent allowing their tissue to be used for research purposes once diagnostic procedures were completed.

Resected human tumor material was immediately placed on ice on in RPMI 1640 (Thermo Fisher Scientific) supplemented with 2.5% fetal bovine serum (FBS) (Sigma-Aldrich) and 1% penicillin-streptomycin (Roche). Tumor samples were processed on a petri dish on ice and manually dissected into small fragments (patient-derived tumor fragments; PDTFs) measuring 1-2 mm³. These fragments were first mixed and then combined into collections of 8-12 PDTFs to ensure representation from different tumor regions. Each collection of PDTFs was frozen in 1 mL of freezing medium (FBS with 10% dimethyl sulfoxide (Sigma-Aldrich)) and cryopreserved in liquid nitrogen until further usage.

### Ex vivo treatment of patient-derived tumor fragments

PDTF cultures were performed following established protocols (*18, 52*). In summary, cryopreserved PDTFs were thawed in a water bath of 37°C until a small ice cube was left. The fragments were then extensively washed with wash medium (Dulbecco’s modified Eagle’s medium (DMEM) supplemented with 10% FCS and 1% penicillin-streptomycin) to remove DMSO, and embedded in an artificial extracellular matrix composed of sodium bicarbonate (1.1%; Sigma-Aldrich), rat-tail collagen I (1 mg/mL; Corning), Matrigel (2 mg/mL; Matrix High Concentration, Phenol Red-Free, BD Biosciences) or Cultrex UltiMatrix (2 mg/mL; reduced growth factor basement membrane extract, R&D systems), and tumor medium [DMEM supplemented with 1 mM sodium pyruvate (Sigma-Aldrich), 1× MEM non-essential AA (Sigma-Aldrich), 2 mM L-glutamine (Thermo Fisher Scientific), 10% FBS, and 1% penicillin-streptomycin]. The embedding procedure started with the addition of 40 µL extracellular matrix to a flat-bottom 96-well plate, which was incubated at 37°C for 15 to 30 min to solidify. During this incubation, cryopreserved PDTFs were thawed as described above and directly placed on top of the matrix layer, after which another 40 µL of soluble matrix was added and incubated at 37°C for 15 to 30 min. To prevent dehydration of the matrix, 20 µL of tumor medium was added to a total volume of 100 µL per well. Ex vivo treatments were prepared by supplementing tumor medium with either anti-PD-1 (Nivolumab, Bristol-Myers Squibb, 10 µg/mL) or anti-CD3 (OKT3, BioLegend, 0.5 µg/mL). For anti-PD-1 perturbations, PDTFs were preincubated for 2 hours at 37°C with either anti-IFNγR1 (50 µg/mL, catalog no. 92101, R&D systems), Lcki (8 μmol/L, CAS no. 213743-31-8, Merck Millipore), αCD4xαPD1 (10 µg/mL, Asher Biotherapeutics) and/or αCD8xαPD1 (10 µg/mL, Asher Biotherapeutics). Importantly, αCD8xαPD1 was designed to bind to CD8β, preventing the targeting of NK cells. Subsequently, PDTF cultures were incubated for 48 hours at 37°C, and shorter times of 8 hours were used for a subset of experiments where indicated.

### PDTF processing for protein analysis

After ex vivo treatment of PDTFs, supernatants for each experimental treatment condition were pooled and frozen at -80°C until cytokine and chemokine analysis was performed. Fragments were pooled per condition, collected into 2 mL digestion medium [RPMI 1640 with 1% penicillin-streptomycin, Pulmozyme (12.6 µg/mL, Roche), and collagenase type IV (1 mg/mL,

Sigma-Aldrich)] on ice, and subsequently digested at 37°C under continuous rotation for 45- 60 min. To obtain single cell suspensions, digested PDTFs were washed with PBS, centrifuged, resuspended in PBS and filtered. Then, cells were transferred to a 96-well U-bottom plate and incubated with human Fc-Receptor Blocking Reagent (eBioscience) and Zombie NIR (BioLegend) for 20 min on ice. Cells were then washed and incubated with the surface-marker antibodies in staining buffer (eBioscience) for 20 min on ice. Afterwards, cells were washed, fixed and permeabilized using Fix/Perm solution (eBioscience) for 30 min at room temperature (RT). Next, fixed cells were washed twice with 1x permeabilization buffer (eBioscience) and incubated with intracellular antibodies in permeabilization buffer for 40 min at RT. Cells were washed twice and resuspended in FACS buffer [PBS supplemented with 2 mM Ethylenediaminetetraacetic acid (EDTA)] for data acquisition on a Symphony A5 (BD Biosciences) or Aurora (Cytek Bio). All antibodies used are displayed in **Table S2**. Data analysis was performed using the FlowJo software (version 10.7).

### Cytokine and chemokine measurements

Supernatants collected from PDTF cultures were thawed on ice for 45-60 minutes and pooled for each experimental condition (using 8–10 replicates per condition). Cytokines, chemokines, and cytotoxic mediators were quantified using the LEGENDplex Human CD8/NK, Human Proinflammatory Cytokine and Human Proinflammatory Chemokine panels (BioLegend). For each tumor, the proinflammatory chemokine panel was combined with either CD8/NK or proinflammatory cytokine panel, and for clustering analysis only the overlapping mediators were used. For downstream analyses, when tumors were pooled in experimental groups, all measured mediators were taken into account. For each assay, 17 µL of sample was analyzed, following the manufacturer’s protocols, and measurements were obtained on a BD LSR Fortessa™ X-20 Cell Analyzer (BD Biosciences). For data analysis, only samples were considered that passed quality control. For this, pairwise comparisons of all untreated and treated samples of a tumor for three highly secreted steady-state parameters, IL-6, IL-8 or CCL2, were performed. Samples with a ≥10-fold deviation of at least one of these mediators from the other conditions of the same tumor were excluded.

### Degranulation and IFN-γ staining

To evaluate degranulation of T cells following αCD4xαPD1 and αCD8xαPD1 incubation, appoximately 200k PBMCs were plated in round-bottom 96-well plates and rested for 1 hour. Cells were then stimulated with αCD3 (OKT3, BioLegend, 5.0 µg/mL) in the presence of αCD4xαPD1 or αCD8xαPD1 (both provided by Asher Biotherapeutics) or anti-PD-1 (Nivolumab, Bristol-Myers Squibb) in varying concentrations, and in the presence of anti- CD107a-FITC (1:50, BioLegend). After 1 hour, GolgiPlug (1:1000) and GolgiStop (1:1500) (both from BD Biosciences) were added, and cultures were incubated for another 12 hours. Samples were then processed for flow cytometry staining, as previously described, and analyzed on an Aurora (Cytek Bio) analyzer.

### PDTF processing for transcriptome analysis

For transcriptome analysis, PDTF cultures were processed as described above until a single cell suspension was obtained. The remaining cell suspension was washed with phosphate-buffered saline (PBS), filtered, and subsequently transferred to 1.5 mL Eppendorf tubes. Cells were then resuspended in 25 µL cold Cell Staining Buffer (Biolegend) with Human TruStain FcX (1:10, Biolegend) and incubated for 10 min on ice. Pooling different experimental conditions for sequencing was made possible by the addition of TotalSeq-C anti-human hashtag antibodies (numbers 1-24, 1 μg/ml final concentration, Biolegend). Next, 25 µL of Cell Staining buffer containing anti-CD45-PerCP-Cy5.5 (1:50, Invitrogen) for the CD45^+^-sorted PDTF experiments, and anti-CD137 (1:50, BV421, biolegend), anti-CD39 (BV786, BD Biosciences) and anti-CD3 (PE, biolegend) for the enrichment experiments. TotalSeq-C antibodies against CD8 (SK1, 1:5000) and CD4 (RPA-T4, 1:2500) were added to these mixes. Cells were subsequently incubated on ice for 25 minutes and then washed three times with 1 mL of staining buffer. They were resuspended in 500 µL of MACS buffer, which consisted of PBS with 0.5% bovine serum albumin (BSA, Sigma) and 2 mM ethylenediaminetetraacetic acid (EDTA, Life Technologies). For the CD45^+^-sorted PDTF experiments, 5 µL aliquots from each sample were counted using AccuCount Blank Particles (13.0-17.9 µm, Spherotech) in order to pool equal cell numbers from each experimental condition labeled with different hashtag antibodies. Dead cells were stained with propidium iodide (PI, Sigma Aldrich, 0.5 µg/ml) just prior to acquisition. Using flow cytometry, live immune cells were counted per 10,000 counting beads. Using a FACSAria Fusion Flow Cytometer (BD Biosciences), CD45^+^ live cells from this mixture were sorted and collected in cold RPMI 1640 medium supplemented with 1% penicillin-streptomycin and 10% human serum. Finally, cells were washed once with cold 1% BSA in PBS and once with 0.04% BSA in PBS, then resuspended in 0.04% BSA at a concentration between 800 and 1200 cells/µL for 10X Genomics scRNA and TCR sequencing. For enrichment of reinvigorated cells, the hashtag staining procedure described above was performed subsequent to the sorting procedure to stain each individual sorted population, after which cells were directly subjected to 10X Genomics scRNAseq.

### Single-Cell RNA and TCR Sequencing

For single cell RNA and TCR sequencing, pooled sorted cells were loaded into each lane of the 10X Chromium instrument, targeting a capture rate of 1,000 to 10,000 individual cells per lane. RNA, TCR, and antibody barcode libraries were created following the manufacturer’s protocol using the Chromium Next GEM Single Cell V(D)J Reagent Kits (10X Genomics). A Novaseq instrument (Illumina) was used to sequence these libraries, with read lengths of 26-28/58-130 for RNA and HTO libraries, and 26-28/92-130 for TCR libraries. Our objective was to reach recovery of 30,000 read pairs per cell for RNA libraries and 5,000 reads for both antibody and TCR libraries.

### Single-cell data analysis pipeline

Sequenced gene expression reads were mapped to the human GRCh38-2020-A reference genome and quantified using Cell Ranger software (10X Genomics, v7.1.0). The filtered gene expression matrix and cite-seq antibody count matrix were imported into Seurat (v5.0.)(*89*) and processed on a per-patient basis, labeling conditions or sorted cell populations with barcoded hashtag oligos (HTOs). Cells lacking a dominant HTO signal or exhibiting multiple dominant HTOs were removed, and datasets were merged. To maintain high data quality, cells were filtered based on mitochondrial RNA content (<15%) and UMI count (>800 or <6000). For the sequenced TCRs, Cell Ranger was used to assemble TCR reads into consensus sequences, excluding cells with multiple TCRβ chains (considered doublets) and identifying T cell clones with matching TCRα and TCRβ sequences.

After preprocessing, transcripts from quality-controlled cells were log10-normalized using Seurat’s NormalizeData() function (*89*) and CITE-seq counts were normalized with a center-log ratio (CLR). Highly variable genes were selected, excluding mitochondrial, ribosomal, long non-coding RNAs, and IG and TCR-V genes. PCA was performed, followed by UMAP and clustering based on a k-nearest neighbors graph with the Louvain algorithm. To improve the accuracy of CD4⁺ and CD8⁺ T cell annotations, cells were filtered based on both gene and protein expression. *CD3E* gene expression was required (> 0.25) to include only T cells. CD4⁺ T cells were defined as having *CD4* gene expression (> 0.25) and *CD8A* and *CD8B* expression

(≤ 0.25), combined with CD4 CITE-seq signal (> 0) and CD8 CITE-seq signal (= 0). CD8⁺ T cells were defined as *CD8A* and *CD8B* expression (> 0.25) and *CD4* expression (≤ 0.25), combined with CD8 CITE-seq signal (> 0) and CD4 CITE-seq signal (= 0). Subsetted CD8^+^ and CD4^+^ T cells were reclustered for detailed subset analysis. Batch and tumor-specific effects were mitigated using the RunHarmony() function(*90*), and UMAP projections were created for cluster identification. Differentially expressed genes between clusters were identified using Seurat’s FindAllMarkers()(*89*) and FindMarkers()(*89*) functions, and gene signatures scores were computed using the AddModuleScore()(*89*) function. Lastly, TCR annotation was performed using the 10X cellranger vdj pipeline, with doublets excluded. The scRepertoire package(*91*) was employed for clonotype assignment and analysis of clonotype dynamics.

### Statistical Analysis

Statistical significance was assessed using two-tailed Wilcoxon test or Friedman tests. Statistical significance thresholds were set as follows: \*\*\*\**P* < 0.0001, \*\*\**P* < 0.001, \*\**P* < 0.01, and \**P* < 0.05. Analyses were conducted using GraphPad (v.10.0.3) or R (v.4.3.3). Due to limited material availability, experiments were generally performed without duplicates unless otherwise noted.

### Supplementary Materials

Supplementary Figure 1: Characterization of αCD4xαPD1, αCD8xαPD1 and αPD-1 effects in patient-derived tumor fragments (PDTFs).

Supplementary Figure 2: Single cell profiling of *PDCD1* positive and negative T cells. Supplementary Figure 3: Short term (8-hour) treatment of PDTFs with αPD-1.

Supplementary Figure 4: Characterization of clusters of intratumoral T cells after ex vivo PDTF culture.

Supplementary Figure 5: Features of dysfunctional subsets within intratumoral CD8^+^ and CD4^+^ T cells.

Supplementary Figure 6: Treatment-induced changes by αPD-1 and αCD3 in intratumoral CD4^+^ and CD8^+^ T cells.

Supplementary Figure 7: Translation-associated gene expression in dysfunctional T cells reinvigorated following αPD-1.

Supplementary Table 1: Patient characteristics of resected tumor samples used for PDTF cultures.

Supplementary Table 2: Human flow cytometry antibodies.

Supplementary Table 3: Genes associated with *PDCD1* expression in intratumoral CD8^+^ and CD4^+^ T cells.

Supplementary Table 4: Published gene signatures references in this study.

## Acknowledgments

We gratefully acknowledge the core facilities at NKI-AVL for their outstanding technical assistance, particularly the Flow Cytometry Facility, including Guido de Roo for assistance with the flow and cell sorting experiments, and the Genomics Core Facility, including Iris de Rink and Arno Velds, for their valuable contributions to the single-cell analyses. Our thanks extend to the Molecular Pathology & Biobanking Core Facility for their help in collecting and processing human tissue samples. Lastly, we want to thank all members of the Thommen laboratory for their constructive discussions. Illustrations in Figure 1E, 2A, and 6B were created using BioRender.com.

## Funding

This research was funded by a Team Science Award from the Melanoma Research Alliance (681127) awarded to D.S.T., in collaboration with Christian Blank and Daniel Peeper, by Oncode Institute through a donation by Mr. H.J.M. Roels (to D.S.T.), by research funding from Asher Biotherapeutics (to D.S.T), and by the Stevin award (to T.N.M.S). We also acknowledge the institutional grant to the NKI from the Dutch Cancer Society (KWF) and the Dutch Ministry of Health, Welfare and Sport.

## Author contributions

Conceptualization: D.S.T., T.N.M.S. Methodology: P.K., N.S., A.M.v.d.L, S.M.C., Y.A.Y., K.D.M., and I.M.D. Investigation: P.K., N.S., A.M.v.d.L, E.R.,R.A.W., T.R., J.M., T.N.M.S. and D.S.T. Visualization: P.K. and D.S.T. Funding acquisition: D.S.T. and T.N.M.S. Project administration: P.K. Supervision: D.S.T., T.N.M.S. Writing – original draft: P.K and D.S.T. Writing – review & editing: all authors.

## Competing Interests

K.D.M., Y.A.Y., and I.M.D. are employed by Asher Biotherapeutics. T.M.R. has received research funding from Bristol Myers Squibb unrelated to the present work. T.N.M.S. serves as an advisor and member of the scientific advisory board and holds shares in Asher Biotherapeutics. T.N.M.S. also advises Allogene Therapeutics, Merus, Neogene Therapeutics, and Scenic Biotech, holds stock in Allogene Therapeutics, Merus, and Scenic Biotech, and is a venture partner at Third Rock Ventures; these roles are unrelated to the present work. D.S.T. received compounds and research funding for this study from Asher Biotherapeutics, and research funding from Bristol Myers Squibb unrelated to the present work.

## Data and Code Availability

Source data will be provided with this publication. The scRNA and TCRseq data generated for this study are deposited in the European Genome-Phenome Archive (EGA) under accession number EGAS50000000881, accessible upon request. For data access or related inquiries, please contact. Additional supporting data are available from the corresponding author upon reasonable request.

## Supplementary Materials

**Figure S1.**
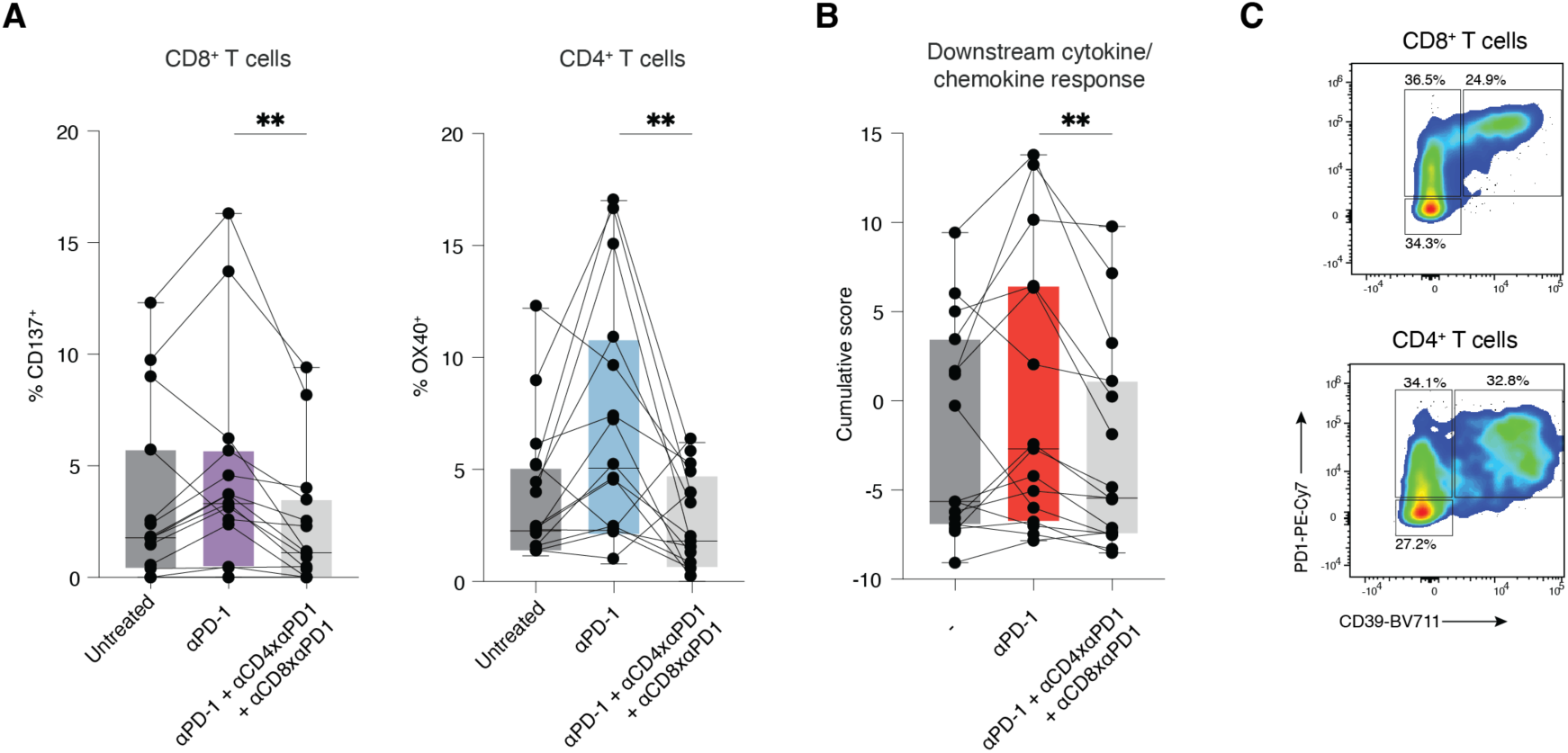
Characterization of αCD4xαPD1, αCD8xαPD1 and αPD-1 effects in patient-derived tumor fragments (PDTFs). (**A**) Quantification of CD137 on intratumoral CD8^+^ T cells and OX40 on intratumoral CD4^+^ T cells from unstimulated, αPD-1, and αPD- 1+αCD4xαPD1+αCD8xαPD1 treated PDTFs, as measured by flow ytometry (n = 15). \*\*\**P* < 0.001, \*\**P* < 0.01 by Friedman test corrected for multiple comparisons. (**B**) Cumulative downstream cytokine/chemokine response score (calculated by adding normalized soluble mediator values for each experimental condition) for each tumor. The soluble mediators considered for this analysis were based on the parameters most discriminative between ex vivo αPD-1 responders and non-responders (*18*), consisting of IFN-γ, IL-10, CXCL9, CXCL10, CXCL11, CXCL5, CCL17, CCL5, CCL4, CCL20 and CXCL1 (n = 15). (**C**) Representative flow cytometry plots displaying the expression of PD-1 and CD39 in intratumoral CD4^+^ and CD8^+^ T cells.

**Figure S2.**
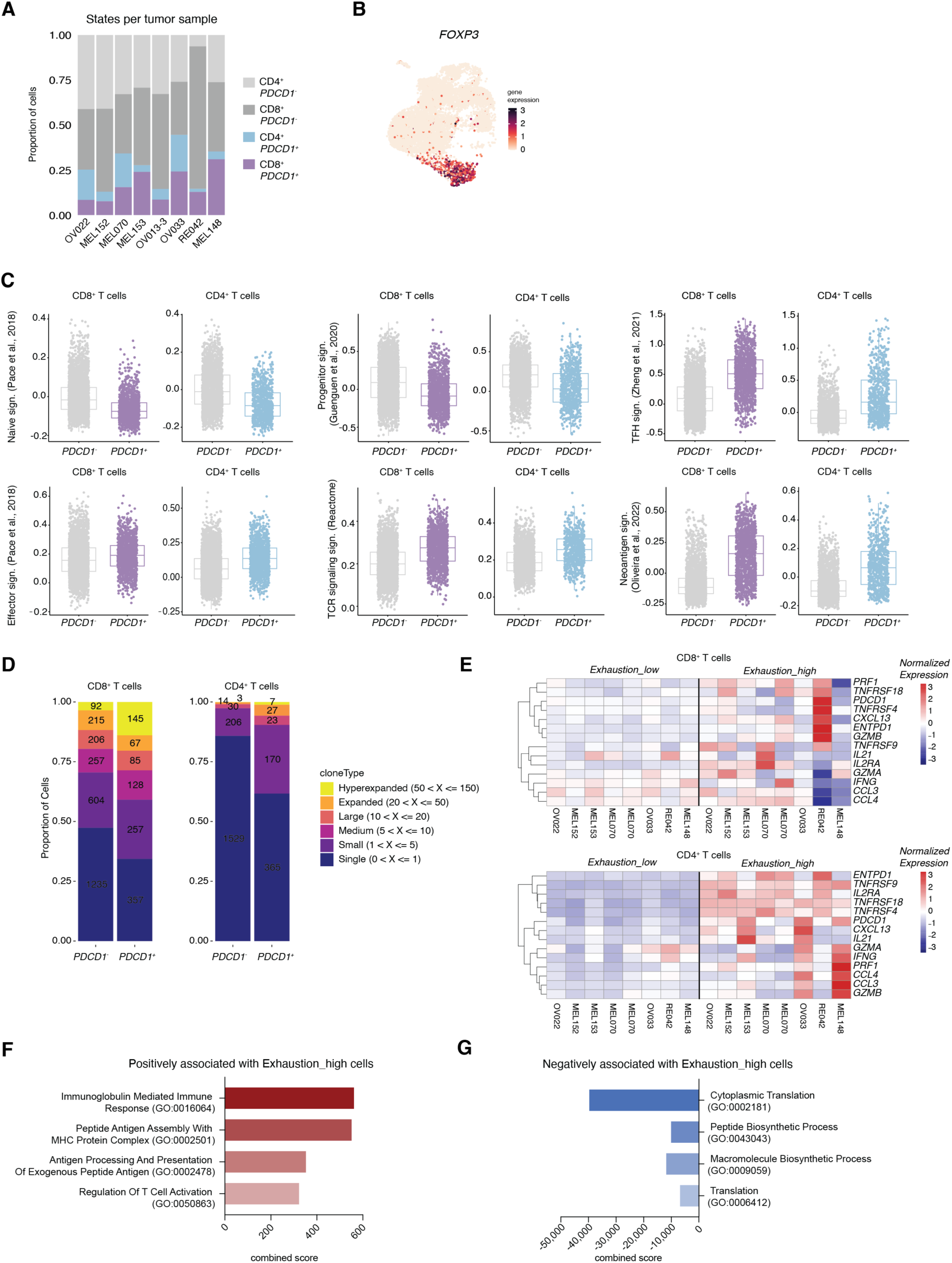
Single cell profiling of *PDCD1* positive and negative T cells. (**A**) Quantification of *PDCD1* positive and negative CD4^+^ and CD8^+^ T cell fractions within total T cells in eight tumors. (**B**) UMAP of intratumoral T cells displaying *FOXP3* gene expression (n = 8). (**C**) Expression of the indicated gene signatures (*30, 58, 62, 63*) in *PDCD1^+^* and *PDCD1^−^* CD8^+^ and CD4^+^ T cells. (**D**) TCR clonotype expansion in *PDCD1*^+^ and *PDCD1*^−^ CD8^+^ and CD4^+^ T cells. TCRs are ranked based on frequency ranges: ‘single’ = found in 1 cell, ‘small’ = in >1 and ≤5 cells, ‘medium’ = in >5 and ≤10 cells, ‘large’ = >10 and ≤20 cells, ‘expanded’ = >20 and ≤50 and hyperexpanded =>50 and ≤150. **(E)** Heatmaps of expression of effector genes in CD8^+^ T cells (top) and CD4^+^ T cells (bottom) separated in Exhaustion_high and Exhaustion_low using an T cell exhaustion signature (*33*) adjusted by removal of effector genes *PDCD1*, *IFNG*, *TNFRSF9* and *CCL3* plotted separately per tumor sample. (**F-G**) Gene Ontology (GO) enrichment analysis of pathways positively (**F**) or negatively (**G**) associated with Exhaustion_high as compared to Exhaustion_low T cells including only cells carrying overlapping TCRs between these groups. X-axis displays a combined score calculated as log(*P*- value) x z-score.

**Figure S3.**
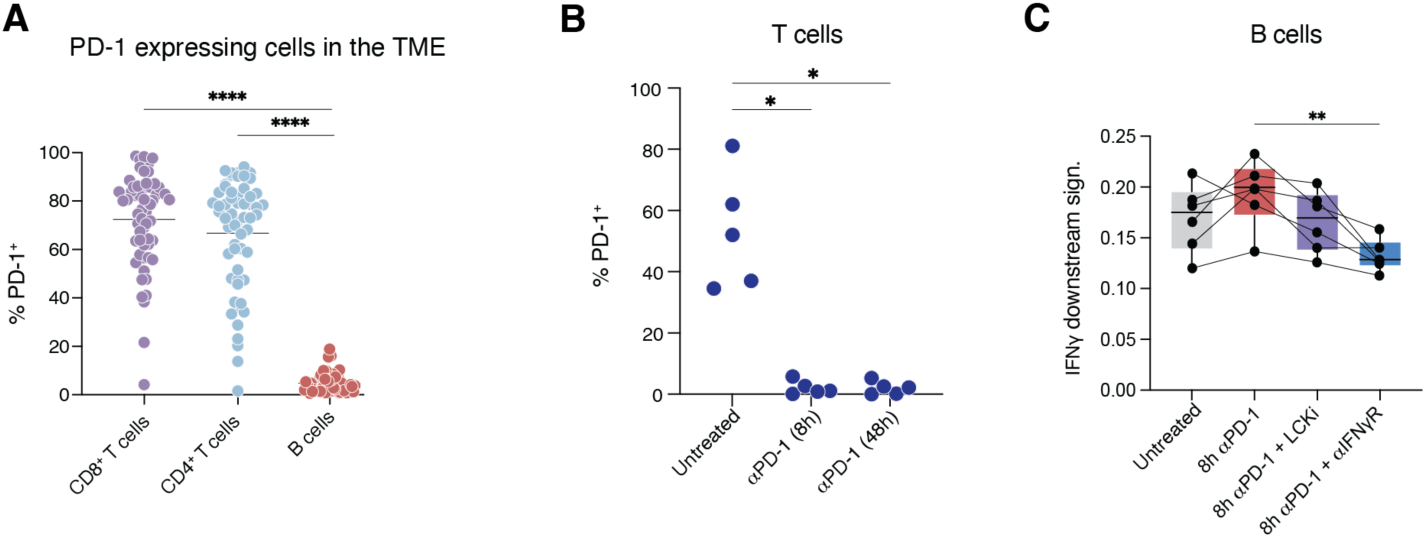
Short term (8-hour) treatment of PDTFs with αPD-1. (**A**) Quantification of the percentage of PD-1 positive intratumoral CD8^+^ T cells, CD4^+^ T cells and B cells in human tumors analyzed directly ex vivo (n = 58). \*\*\*\**P* < 0.0001 by the Friedman test corrected for multiple comparisons. (**B**) PD-1 staining, as measured by flow cytometry in CD3^+^ T cells in PDTFs that were left untreated or treated with 8 and 48 hours αPD-1, respectively, ex vivo (n = 5). \**P* < 0.05 by Friedman test corrected for multiple comparisons. (**C**) IFN-γ response gene signature (*36*) in B cells separated per tumor for PDTFs treated ex vivo for 8 hours. \*\**P* < 0.01 by Friedman test corrected for multiple comparisons.

**Figure S4.**
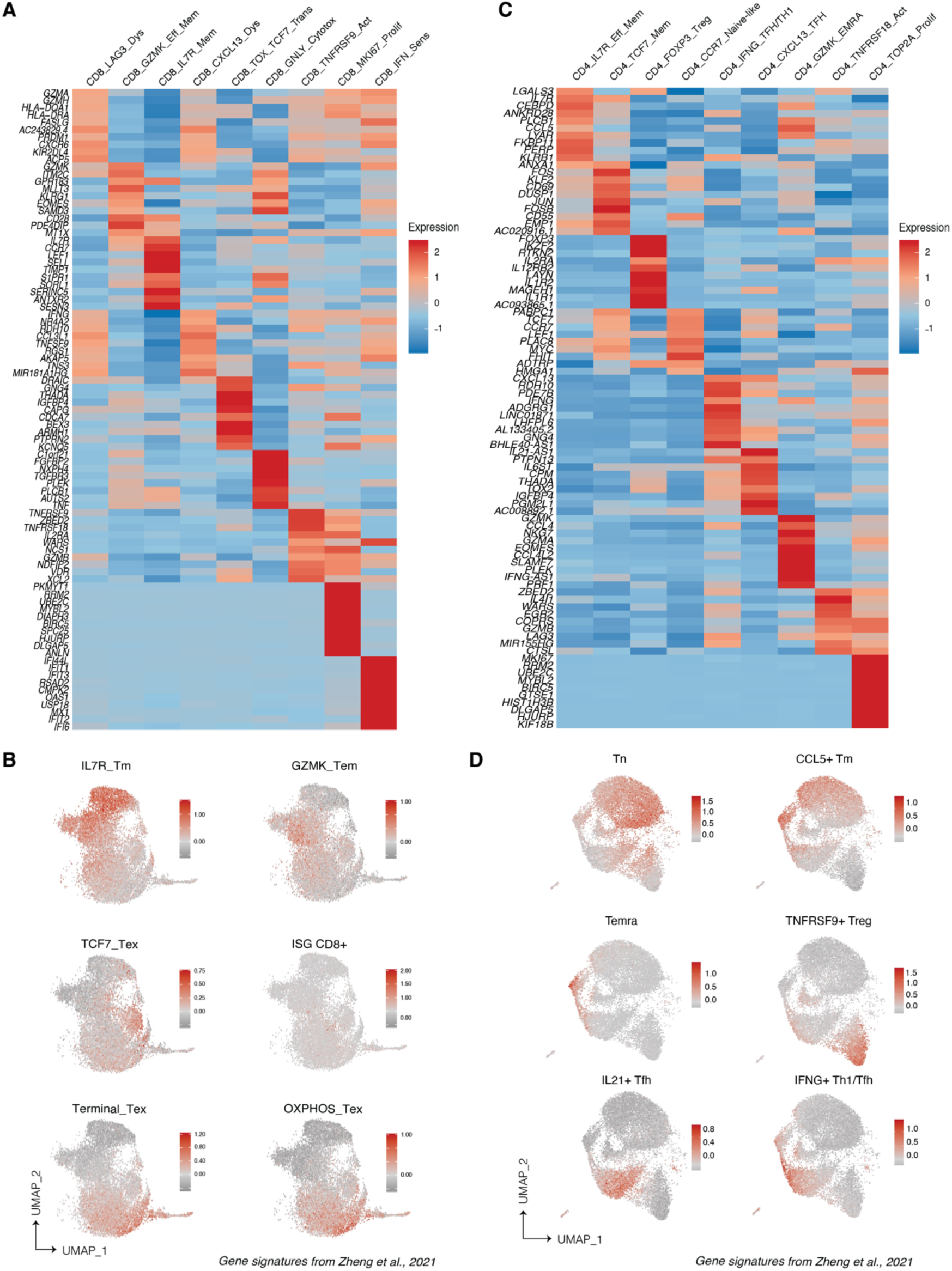
Characterization of clusters of intratumoral T cells after ex vivo PDTF culture. **(A)** Heatmap of normalized expression of top10 differentially expressed genes per CD8^+^ T cell cluster (Wilcoxon rank sum test). (**B**) UMAPs of intratumoral CD8^+^ T cells displaying expression of indicated gene signatures (n = 6 tumors) (*58*). (**C**) Heatmap of normalized expression of top10 differentially expressed genes per CD4^+^ T cell cluster (Wilcoxon rank sum test). (**D**) UMAPs of intratumoral CD4^+^ T cells displaying expression of indicated gene signatures (n = 6 tumors) (*58*).

**Figure S5.**
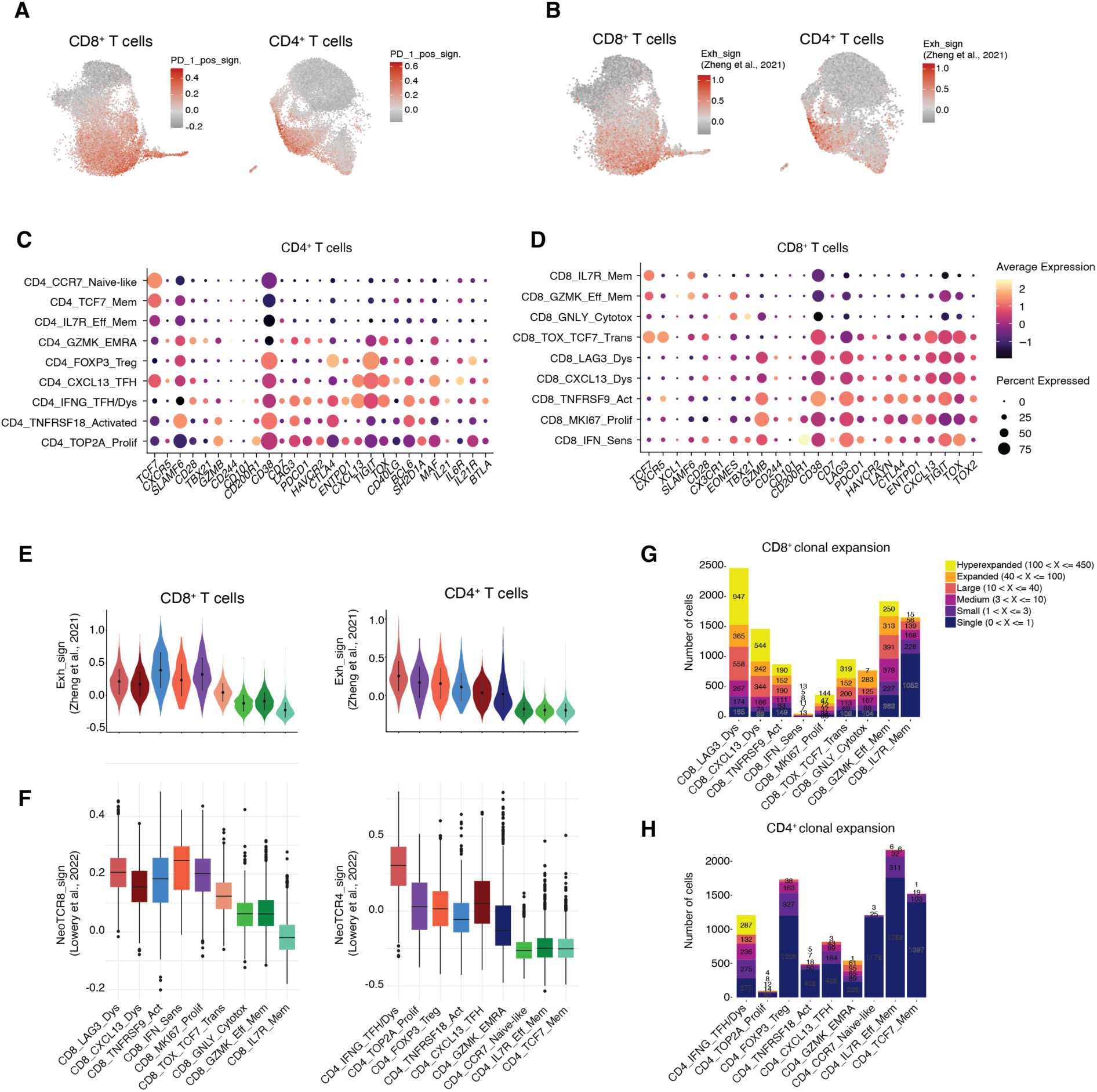
Features of dysfunctional subsets within intratumoral CD8^+^ and CD4^+^ T cells. **(A)** UMAPs displaying enrichment of the gene signature of *PDCD1*^+^ CD8^+^ and CD4^+^ T cells identified in Figure 3D. (**B**) UMAPs displaying enrichment of an exhaustion gene signature (*59*) in intratumoral CD8^+^ and CD4^+^ T cells. (**C-D**) Bubble plots of expressed genes associated with precursor-exhausted and exhausted T cells in chronic viral infections (*6, 40, 67*) in intratumoral CD4^+^ T cells (**E**) and CD8^+^ T cells (**F**). (**E**) Violin plot of the exhaustion gene signature (*59*) per cluster in intratumoral CD8^+^ and CD4^+^ T cells. (**F**) Boxplot of the NeoTCR8 and NeoTCR4 signature scores (*29*) per cluster in intratumoral CD8^+^ and CD4^+^ T cells. (**G-H**) TCR clonotype expansion of CD4^+^ T cells (**F**) and CD8^+^ T cells (**H**) per cluster from PDTFs (n = 6).

**Figure S6.**
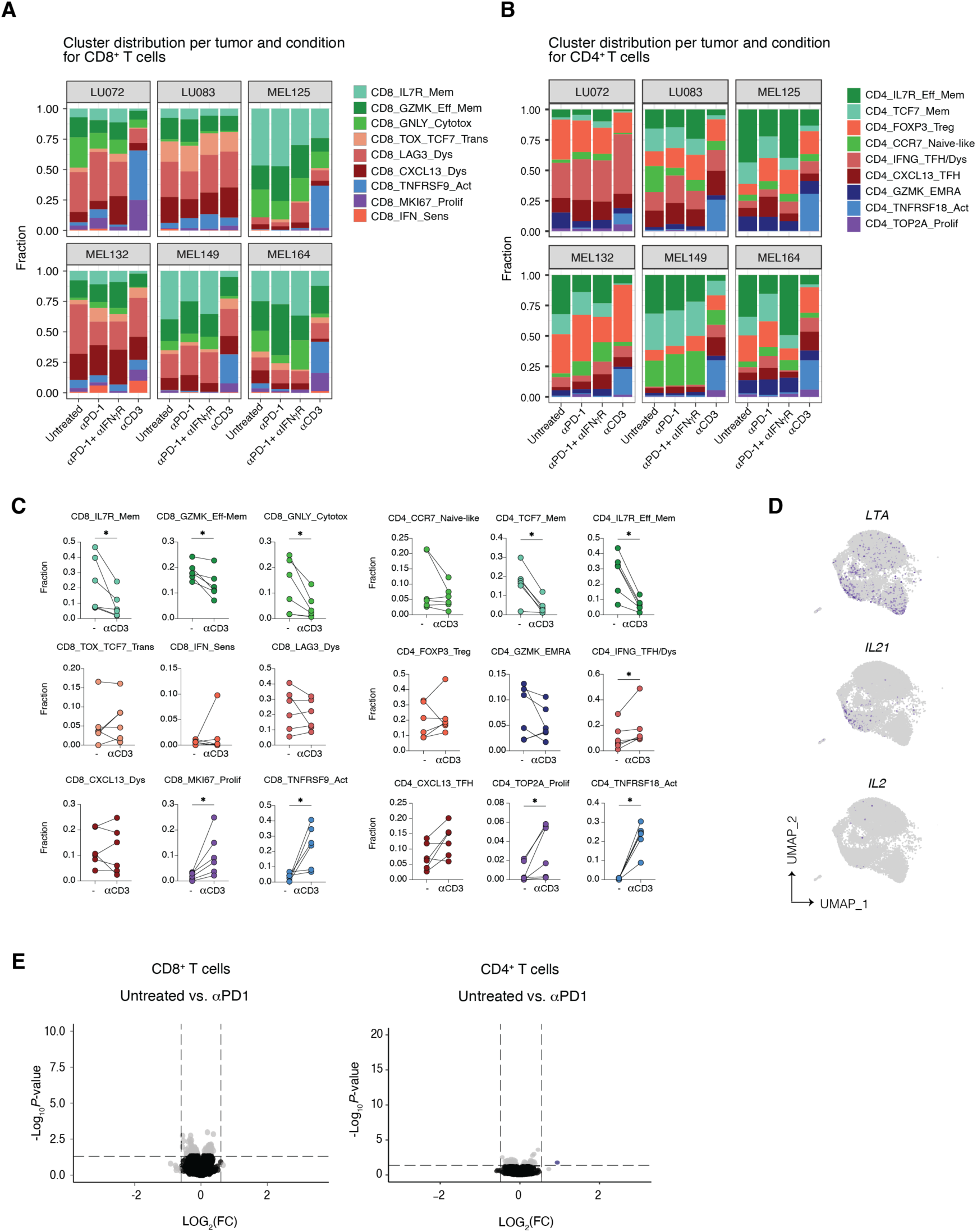
Treatment-induced changes by αPD-1 and αCD3 in intratumoral CD8^+^ and CD4^+^ T cells. (**A+B**) Bar graphs of cluster fractions of CD8^+^ (**A**) and CD4^+^ (**B**) T cell states derived from untreated, αPD-1, αPD-1 + αIFNγR, and αCD3-treated conditions (48 hours). (**C**) Connected dot-plots of cluster fractions of CD8^+^ and CD4^+^ T cell clusters from the untreated and αCD3-treated conditions (n = 6). \**P* < 0.05 by Wilcoxon matched-pairs test. (**D**) UMAPs of intratumoral CD4^+^ T cells displaying expression of the cytokines *LTA, IL21* and *IL2*. (**E**) Differential gene expression analysis (pseudo-bulk, n = 6) between the untreated and short-term αPD-1-treated conditions (8 hours) in CD8^+^ (left) and CD4^+^ T cells (right).

**Figure S7.**
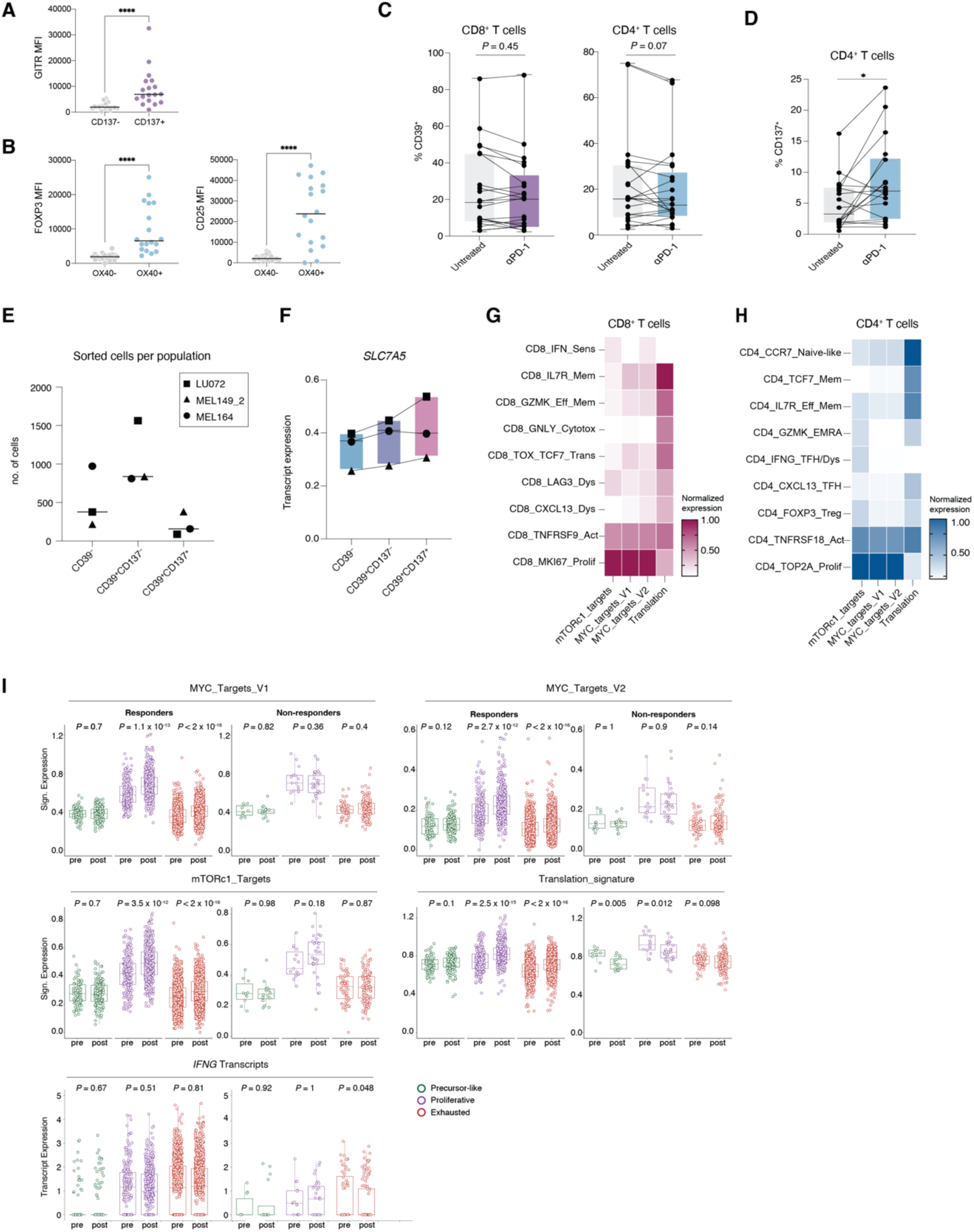
Translation-associated gene expression in dysfunctional T cells reinvigorated following αPD-1. (**A**) Difference in GITR expression measured by mean fluorescence intensity (MFI) in CD137^−^ and CD137^+^ CD8^+^ T cells from αPD-1-treated PDTFs (n = 18). \*\*\*\**P* < 0.0001 by Wilcoxon matched-pairs test. (**B**) Difference in FOXP3 and CD25 expression measured by MFI in OX40^−^ and OX40^+^ CD4^+^ T cells from αPD-1-treated PDTFs (n = 18). \*\*\*\**P* < 0.0001 by Wilcoxon matched-pairs test. (**C**) Quantification of CD39 on total intratumoral CD8^+^ and CD4^+^ T cells in untreated and αPD-1-treated PDTFs, as measured by flow cytometry (n = 18). *P-*value by Wilcoxon matched-pairs test. (**D**) Quantification of CD137 on total intratumoral CD4^+^ T cells in untreated and αPD-1-treated PDTFs, as measured by flow cytometry (n = 18). \**P* < 0.05 by Wilcoxon matched-pairs test. (**E**) Number of sorted cells in the indicated cell populations for three tumor samples. (**F**) Transcript expression of the solute carrier *SLC7A5* in the indicated cell populations for three tumor samples. (**G-H**) Heatmap of min-max normalized expression of mTORc1_targets, MYC_targets_V1, MYC_targets_V2, and translation (Reactome) gene signatures in CD8⁺ T cell clusters (**G**) and CD4^+^ T cell clusters (**H**). (**I**) Gene signatures scores of mTORc1_targets, MYC_targets_V1, MYC_targets_V2, and translation (Reactome) signatures in precursor-like, proliferative and exhausted CD8^+^ T cells in pre- and on-treatment biopsies from breast cancer patients treated with anti-PD-1, separately shown for responders and non-responders. Data was re-analyzed from Bassez et al., 2021 (*74*) and annotated according to Liu et al., 2022 (*56*). *P-*values by Wilcoxon matched-pairs test.

